# Time-Varying Dynamic Causal Modelling for Sequential Responses: Neural Mechanisms of Slow Cortical Potentials, Preparation, Planning and Beyond

**DOI:** 10.64898/2026.03.24.714008

**Authors:** Andrew D Levy, Peter Zeidman, Karl Friston

## Abstract

Cognitive processes such as decision-making, working memory, and motor planning operate across a hierarchy of timescales, manifesting as rapid neural transients alongside slower physiological mechanisms like short-term plasticity. Conventional Dynamic Causal Modelling (DCM) limits our ability to study these dynamics by assuming stationary parameters, whilst recent time-varying approaches often rely on segmenting data into epochs. This segmentation artificially resets neural states between windows, fundamentally obscuring the continuous hysteresis essential to sequential processing. To address this limitation, we introduce DCM for Sequential Responses (DCM-SR), a generative framework that embeds parameter evolution directly within the first-level model whilst employing a continuous state-space formulation that removes the requirement for epoching. This approach generalises non-stationarity to all neural mass parameters, including synaptic gains and time constants, modelling them as piecewise smooth trajectories that evolve alongside continuous neural states. Consequently, the model explicitly captures two distinct forms of temporal memory: transient history dependence, where responses are shaped by the carryover effect of recent perturbations, and path dependence, where the system’s trajectory through parameter space determines its responsiveness. The framework accommodates both exogenous, stimulus-locked transitions and endogenous, autonomous state changes, permitting inference on both external perturbations and internal drivers of network evolution. Simulations establish the model’s face validity, demonstrating robust parameter recovery and conservative model selection that accurately discriminates between genuine parameter evolution and spurious complexity. We applied the framework to empirical data from an auditory go/no-go task, modelling a full sequence of cognitive phases from initial cue processing and anticipation through to motor preparation and execution. This analysis established construct validity by resolving the biophysical generators of the contingent negative variation, attributing this slow potential to sustained thalamocortical drive and deep-layer hyperpolarisation rather than superficial-layer activity. Furthermore, the model captured trial-specific modulations of the hyperdirect pathway during motor inhibition, tracking the dynamic interplay between prefrontal executive control and basal ganglia gating. DCM-SR offers the first principled approach to decomposing compound signals such as slow cortical potentials into evolving synaptic mechanisms and continuous state trajectories, and provides a necessary bridge for investigating the biophysical implementation of extended cognitive phenomena including evidence accumulation and physiological hysteresis.

## 1 Introduction

The brain is situated within a body and world characterised by statistical regularities that unfold over time. A gust of wind on a cloudy day during the winter (or spring) months. The stress placed on a phoneme within a word during a friendly (or hostile) conversation. A heartbeat on a run in the final (or first) mile of a marathon. As such, percepts do not arise in isolation or without prior context, but as temporally structured events whose dependencies inform interpretation and constrain expectations. Consequently, an agent navigating its environment must continuously integrate information across time, updating predictions about what comes next while maintaining representations of what came before, encompassing not only external events but the evolving states of its own body. This continuous integration operates across a hierarchy of temporal scales (Kiebel, Daunizeau, and Friston 2008). Where rapid sensory encoding unfolds over milliseconds, perceptual inference operates across hundreds of milliseconds, working memory maintains information over seconds to minutes, and learning consolidates statistical regularities over hours to days. The brain’s hierarchical temporal architecture thus mirrors the nested temporal structure of the environments within which it evolved, enabling perception and action across the multiple timescales that characterise embodied existence.

Investigating how the brain integrates information across nested timescales requires sequential paradigms that replicate the hierarchical temporal structure characteristic of real-world environments. Consider some classic examples: Posner cueing tasks probe anticipatory attention by invoking preparatory states through a directional cue and subsequent delay prior to the target stimulus (Petersen and Posner 2012); N-back tasks engage sustained working memory by requiring participants to detect matches against stimuli presented N trials earlier (Owen et al. 2005); oddball paradigms access the brain’s sensitivity to statistical structure by presenting sequences of repeating stimuli with occasional deviants (Huettel and McCarthy 2004). Sequential paradigms implicitly operate across multiple timescales, ranging from hundreds of milliseconds to minutes. They typically comprise different stimulus types separated by variable delays, and in more sophisticated protocols, statistical contingencies that unfold across trials. As a consequence, the neural dynamics they invoke evolve continuously between events, shaped by the hierarchical temporal architecture of the brain itself. Understanding how the brain processes such sequences requires generative models capable of capturing this continuous evolution, explicitly representing how neural states and connectivity parameters evolve across time to shape responses through the accumulated influence of preceding context, ongoing preparatory states, and neuromodulatory mechanisms.

Dynamic Causal Modelling (DCM) for electrophysiological data has proven invaluable for characterising effective connectivity underlying evoked responses to discrete experimental events (David et al. 2006). Conventional DCM implementations assume stationary parameters that remain fixed across experimental epochs, an approximation that holds well for short periods spanning hundreds of milliseconds. However, the paradigms and temporal scales discussed above challenge this assumption. Between successive events, electrophysiological observations reveal continuously evolving dynamics: slow cortical potentials such as the contingent negative variation (CNV) and readiness potential ramp over seconds (Walter et al. 1964; Kornhuber and Deecke 1965), sustained activity patterns track working memory maintenance (Mongillo, Rumpel, and Loewenstein 2018; Stokes 2015), and ramping signals reflect evidence accumulation (Gold and Shadlen 2007). These data features index cognitive and neural processes operating at temporal scales where synaptic mechanisms such as short-term plasticity are inherently dynamic, precluding the assumption of stationary synaptic efficacy parameters. Crucially, accounting for these slow-evolving dynamical processes remains tractable due to a separation of timescales, where neural membrane potentials fluctuate on millisecond timescales whilst synaptic efficacies and neuronal gain evolve over hundreds of milliseconds to seconds (Medrano, Friston, and Zeidman 2024). This separation, formalised through geometric singular perturbation theory (Fenichel 1979), permits decomposition into fast neuronal responses and slowly evolving parameters. Where the fast neuronal states rapidly equilibrate to a manifold determined by the current slow parameters, whilst the slow parameters evolve on a timescale over which the fast states remain approximately stationary. As fast states equilibrate rapidly relative to parameter evolution, the system dynamics can be tractably decomposed into fast transients that quickly settle onto a manifold defined by the current parameter values, and slow parameter evolution that causes this manifold itself to drift over time.

Time-varying DCM approaches exploiting timescale separation have been introduced. One incorporates a Hidden Markov Model to explain time-varying synaptic efficacies, modelling itinerant reconfigurations characteristic of resting-state activity through heteroclinic orbit dynamics and probabilistic transitions between discrete metastable connectivity states (Zarghami and Friston 2020). Whilst theoretically grounded and computationally tractable, the discrete-state assumption cannot capture scenarios where connectivity varies continuously or when transitions are deterministically triggered rather than stochastically governed. Another approach uses an adiabatic approximation with hierarchical parametric empirical Bayes (Jafarian et al., 2021), fitting separate DCMs to recordings partitioned into short epochs, then estimating smooth parameter trajectories post-hoc from posterior estimates using a GLM-based approach constrained with basis sets. Whilst using posteriors from the previous window as priors for subsequent windows attempts to eschew local minima, this does not guarantee convergence to the same local minimum across windows. Parameter trajectories may therefore reflect discontinuous jumps between solutions rather than smooth trajectories through coherent parameter space. A variant addresses this by incorporating temporal variation directly at the first level, thereby avoiding entrapment in different local minima across epochs. This approach models time-varying modulatory connectivity as weighted sums of temporal basis functions evaluated during forward integration, estimating all parameters jointly within a single optimisation (Medrano et al., 2024). This unified framework substantially improves computational efficiency and parameter identifiability, representing the current state-of-the-art for continuous time-varying task-evoked DCM. However, two fundamental constraints limit this approach’s applicability to sequential paradigms. First, time-variation is restricted to modulatory connectivity, maintaining time-invariant baseline connectivity, synaptic gains, time constants, and transmission delays; inappropriate when physiological state changes require comprehensive parameter evolution. Second, it inherits from conventional DCM the requirement for epoched neural responses aligned to discrete events. Whilst this approach models smooth parameter trajectories, the neural data remain segmented into discrete time-locked responses, precluding characterisation of hysteresis and continuous dynamics of processes like sustained attention and motor preparation that unfold across experimental structure.

DCM for Sequential Responses (DCM-SR) evolves from basis-DCM by extending parameter trajectories within the first-level generative model to all key parameters of neural mass models. The approach represents parameter evolution through piecewise smooth trajectories defined at discrete sequential events, whilst neural states propagate continuously through the time-varying dynamical system. Like conventional DCM for MEG/EEG, which has proven valuable for disambiguating neural generators underlying diverse electrophysiological markers (Garrido et al. 2008), DCM-SR decomposes compound signals such as slow cortical potentials in terms of slowly evolving effective connectivity and synaptic mechanisms, manifesting over continuous neural states that bridge multiple events without requiring windowing or epoching. For instance, slow cortical potentials reflect mixtures of cognitive processes, anticipatory states, memory operations, and decision variables, exhibiting well-replicated but non-specific mappings to behaviour and neuronal processes: CNV amplitude scales with preparation and temporal certainty yet varies with arousal and motivation (Brunia and van Boxtel 2001), ramping activity could reflect evidence accumulation, urgency signals, or reward discounting (Churchland, Kiani, and Shadlen 2008; Hawkins et al. 2015), and theta power increases during both encoding and retrieval (Hanslmayr, Staudigl, and Fellner 2012). By explicitly modelling time-varying parameters alongside continuous neural states, DCM-SR provides a principled approach to resolving these ambiguities, moving beyond phenomenological characterisation to mechanistic inference about the dynamic reconfigurations of effective connectivity generating observed responses.

The present paper introduces DCM-SR and establishes its validity through simulation studies and empirical applications. We first detail the mathematical framework and computational implementation, showing how time-varying parameters are integrated within the generic DCM architecture (van Wijk et al. 2018). Simulation studies then demonstrate appropriate model selection and parameter recovery under controlled conditions, including disambiguation of exogenous inputs from endogenous parameter evolution. We subsequently illustrate the framework’s application to an empirical auditory go/no-go dataset, examining how effective connectivity evolves across extended cue-to-target intervals spanning seconds. Together, these components establish DCM-SR as a principled and computationally tractable approach to modelling continuous neural dynamics and time-varying parameters.

## 2 Methods

This section details the implementation and validation of DCM-SR. We begin by specifying the theoretical foundations and technical specification, showing how time-varying parameters are represented through piecewise smooth trajectories indexed by discrete sequential events. We then establish validity through simulation studies demonstrating appropriate model selection and parameter recovery, including disambiguation of exogenous driving inputs from endogenous parameter evolution. Finally, we illustrate the framework’s application to an empirical auditory go/no-go dataset featuring extended cue-to-target intervals, examining how effective connectivity evolves across seconds whilst neural states propagate continuously between events.

### 2.1 DCM for Sequential Responses

DCM-SR follows the principle established by Basis-DCM of embedding parameter evolution directly within the first-level generative model, but generalises this to encompass all key neural mass parameters rather than restricting non-stationarity to modulatory connectivity alone. The approach represents parameter trajectories as piecewise smooth functions, with parameters occupying distinct set-points separated by transition boundaries. These boundaries may be specified as coincident with exogenous stimuli, at approximated timepoints corresponding to endogenous cognitive processes, or synchronised to physiological processes.

This implementation appeals to an adiabatic approximation whereby parameters evolve smoothly on timescales substantially slower than neural membrane dynamics, permitting fast neuronal responses to equilibrate quasi-statically on the manifold defined by slowly varying parameters. The approach distinguishes itself from prior time-varying DCM formulations in two key respects. First, it models continuous neural state propagation across the entire observation period without epoch segmentation. Second, it permits substantial network reconfiguration between transitions, with each transition end-point independently estimated to accommodate drastic connectivity reorganisation.

#### 2.1.1 Technical Details

We begin by clarifying terminology that delineates both the temporal structure of sequential experimental paradigms and the corresponding elements of the generative model. An epoch denotes the entire observation period subject to model inversion, typically spanning several seconds and encompassing multiple experimental events. A paradigm is termed sequential when this epoch contains events presented in a specified and consistent order across trials. An event refers to a discrete occurrence within this sequence that perturbs or modulates the neural system.

The model architecture distinguishes between fast and slow dynamics through hierarchical organisation of states. Neural states comprise the fast variables that evolve on millisecond timescales according to neural mass differential equations. Parameters govern the dynamics of these neural states. Whilst conventional DCM holds parameters fixed throughout the epoch, the approach developed here permits them to vary smoothly across time as parameter states, evolving continuously on substantially longer timescales of hundreds of milliseconds to seconds.

We adopt dynamical systems terminology to characterise the behaviour of our time-varying parameter model. Specifically, we refer to the system’s ‘regime’ to denote a particular parameter configuration and use ‘regime shifts’ to describe rapid transitions between qualitatively distinct configurations, whilst ‘parameter drift’ describes gradual changes that occur within a stable regime. It is important to emphasise that we employ these concepts as an interpretive framework rather than through explicit modelling of the underlying dynamical structure. Our model does not formally represent bifurcation diagrams, attractor basins, or stability landscapes. Instead, we use a functional approximation to capture the temporal characteristics of parameter trajectories. This approach allows us to distinguish between different timescales of change: rapid transitions that we interpret as regime shifts versus slower, continuous evolution that we interpret as drift. The functional approximation thus provides a tractable method for describing these dynamical phenomena whilst remaining agnostic about the precise mechanistic structure that would generate such behaviour in a fully specified dynamical system.

We adopt dynamical systems terminology to characterise the behaviour of our time-varying parameter model. Specifically, we refer to the system’s ‘regime’ to denote a particular parameter configuration, and use ‘regime shifts’ to describe rapid transitions between distinct regimes, whilst ‘parameter drift’ describes gradual changes within a single regime. It is important to emphasise that we employ these concepts as an interpretive framework rather than through explicit modelling of attractor dynamics. Our model does not formally represent bifurcation structure, attractor basins, or stability landscapes. Instead, we use a functional approximation to capture the temporal characteristics of parameter trajectories. This approximation allows us to distinguish between timescales: rapid transitions that we interpret as regime shifts versus slow, continuous changes that we interpret as drift.

##### 2.1.1.1 Trajectory function

DCM-SR extends the standard DCM state-space formulation to accommodate time-varying parameters through piecewise smooth interpolation. The system is described by:

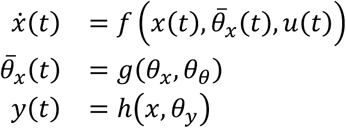

where *ẋ*(*t*) governs the fast neural state dynamics (e.g. membrane potentials and currents) evolving on millisecond timescales, *θ̄*_*x*_(*t*) represents time-varying parameter states (e.g. membrane potentials and currents) evolving on substantially slower timescales, and *y*(*t*) is the observed neural activity. The function *g*(*θ*_*x*_, *θ*_*θ*_) specifies an algebraic interpolation procedure that reconstructs continuous parameter trajectories from discrete regime-specific values using trajectory hyperparameters, *θ*_*θ*_, that govern transition onset and dispersion. Crucially, *θ̄*_*x*_(*t*) is prescribed through interpolation rather than evolved through coupled differential equations, reflecting an adiabatic approximation where slow parameter dynamics are assumed separable from fast neural dynamics.

The interpolation function constructs parameter trajectories as weighted sums of discrete parameter values:

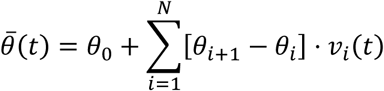

where *N* denotes the number of transitions, *θ_i_* represents the discrete parameter value estimated for regime *i* (Figure 1A), and *v_i_*(*t*) is the trajectory function that weights the contribution of each discrete parameter over time (Figure 1B). This formulation expresses *θ*(*t*) as an initial value plus cumulative increments weighted by trajectory functions that smoothly interpolate between successive regimes (Figure 1C). The trajectory function *v*(*t*) can be parameterised in multiple ways depending on hypothesised parameter dynamics. For sustained state changes, monotonic functions such as sigmoids provide appropriate representations. For transient modulations, functions such as B-splines or Gaussian kernels may be more suitable.

**Figure 1.**
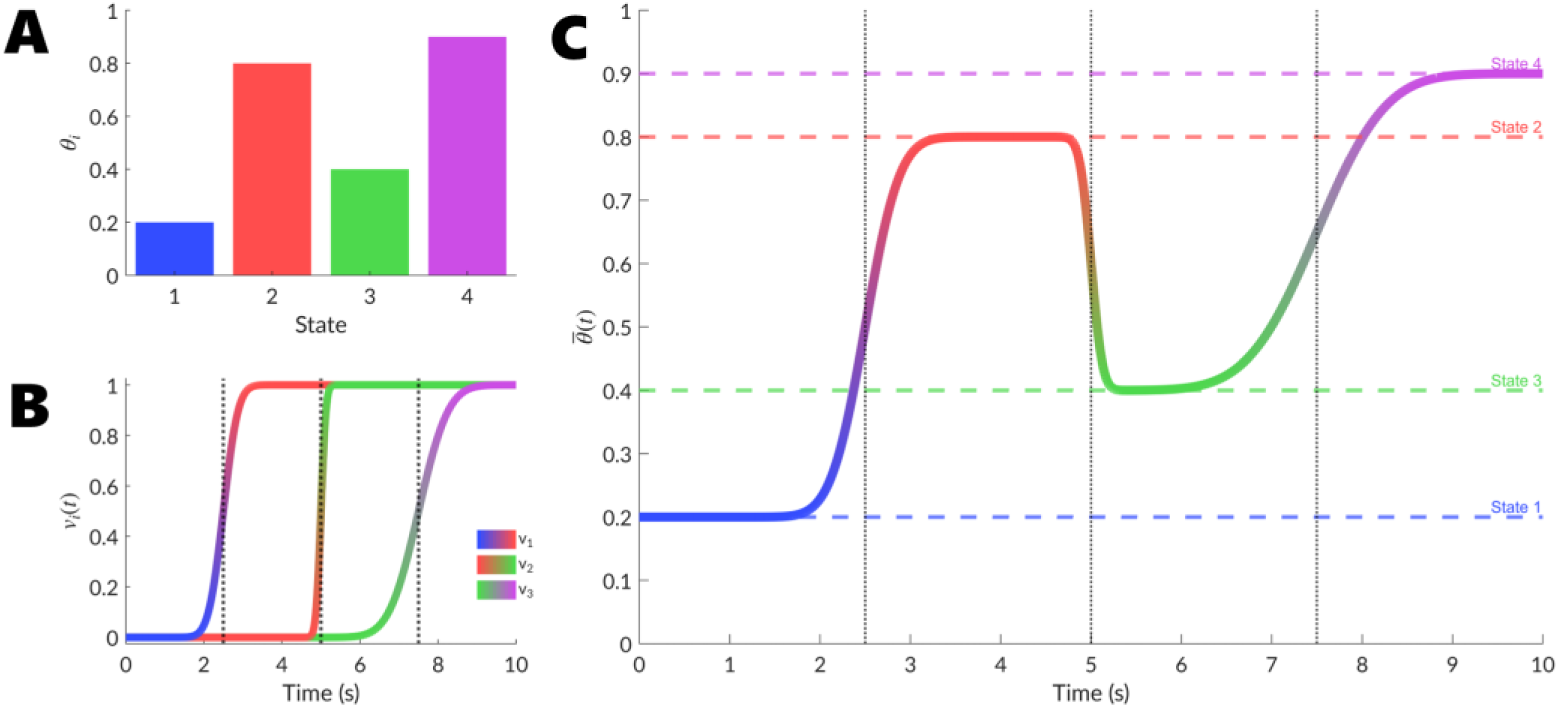
Construction of parameter trajectories through sequential state transitions. This schematic illustrates how DCM-SR generates smooth parameter trajectories across multiple experimental events through interpolation between discrete states. (A) Four discrete parameter states (*θ*_*i*_) are defined at specific timepoints, each characterising the network configuration during distinct experimental phases. (B) Smooth trajectories *v*_*i*_(*t*) for individual parameters are constructed by interpolating between adjacent states using sigmoid transition functions, ensuring continuous evolution whilst preserving differentiability. Vertical dashed lines denote the centres of transition periods between states, with parameter values evolving progressively rather than changing instantaneously. (C) Complete parameter trajectories *θ̄*(*t*) spanning the entire observation period, demonstrating how multiple parameters evolve independently across the experimental structure. Each trajectory smoothly transitions between state-specific values (indicated by horizontal dashed lines), generating time-varying connectivity and neural mass parameters that capture endogenous network reconfiguration during protracted cognitive processes.

The trajectory functions have been implemented such that any neural mass parameter may be specified as time-varying or held constant during model construction. This includes baseline extrinsic connections (A), baseline intrinsic connections (G), modulated driving connections (B), synaptic time constants (T), propagation delays (D), sigmoid activation function slopes (S), and receptor densities (H). This flexibility permits substantial expressivity in capturing parameter dynamics across cognitive states, behavioural epochs, or physiological cycles.

In the present implementation, we employ sigmoid trajectory functions using cumulative Gaussian error functions:

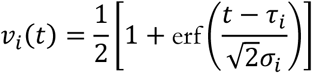

where *τi* specifies the temporal centre of a transition and *σi* determines the transition width, both estimated with priors typically centred on experimental input timing and duration. The width parameter is log-transformed to enforce positivity:

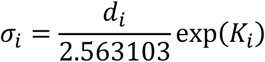

where *di* represents the nominal transition duration, and *Ki* is the estimated log-scale parameter.

Whilst this piecewise sigmoid formulation does not explicitly model underlying bifurcation structure or attractor dynamics, it provides a phenomenological approximation capturing the essential temporal characteristics of these processes. The slope parameter *σ*_*i*_ serves as a proxy for the rate at which the system’s dynamical landscape is reconfigured, with the distinction between drift and shift reflected in the timescale over which parameter values evolve. Large *σ*_*i*_ values produce gentle slopes representing gradual parameter drift within the same attractor basin, capturing processes such as progressive synaptic strengthening, attentional engagement, or sustained neuromodulatory changes that unfold over hundreds of milliseconds. Conversely, small *σ*_*i*_ values produce steep slopes approximating rapid regime shifts, where swift reconfigurations of the dynamical landscape occur over tens of milliseconds. These are characteristic of transitions between attractor basins, as seen in perceptual switches, motor execution onset, or abrupt attentional network reconfigurations.

A crucial assumption is that trajectory functions *v*_*i*_(*t*) are global across all time-varying parameters, sharing identical transition timings *τ*_*i*_ and widths *σ*_*i*_, whilst the discrete parameter values *θ*_*i*_ remain unique to each parameter type. When the system transitions between regimes, all time-varying parameters undergo coordinated reconfigurations governed by the same temporal profile, each evolving to its own estimated value. This design assumes that physiological state transitions constitute global network reconfigurations rather than independent parameter fluctuations, substantially reducing dimensionality whilst preserving the flexibility to capture heterogeneous changes across different connection types.

##### 2.1.1.2 Continuous neural states

Current time-varying DCM approaches epoch data around individual events, treating each as an isolated response. Whilst such approaches can capture smooth parameter changes across epochs, they fundamentally obscure history-dependent carryover by resetting neural states to baseline at each epoch boundary. DCM-SR’s capacity to model continuous neural state evolution enables characterisation of two distinct forms of temporal memory that shape sequential responses. The first is history dependence, where responses reflect transiently altered neural states inherited from preceding events. The second is path dependence or hysteresis, where sustained parameter reconfigurations fundamentally alter the system’s response profile.

###### History-dependent dynamics through neural states

When stimuli are presented in rapid succession, the response to a second stimulus depends not only on its own properties but also on the neural state inherited from the first stimulus. This history dependence arises from multiple fast cellular mechanisms including voltage-gated ion channel kinetics, rectifying currents, rapid synaptic depression through vesicle depletion, and calcium-dependent adaptation currents. Neural mass models implicitly capture these processes through lumped parameters such as synaptic time constants, gains, and recurrent connectivity that aggregate underlying biophysical mechanisms. These intrinsic nonlinearities mean that the response to successive stimuli cannot be predicted by linear summation of isolated responses. Figure 2 (red traces) demonstrates this through simulated responses to two sequential inputs separated by 300ms, where the observed response with stationary parameters (solid line) deviates substantially from the linear prediction (dashed line) obtained by summing isolated stimulus responses. This deviation reveals the nonlinear carryover effects inherent in neural mass dynamics, a form of short-term memory where recent perturbations transiently shift the system’s operating point through altered neural states that subsequently relax back to equilibrium.

**Figure 2.**
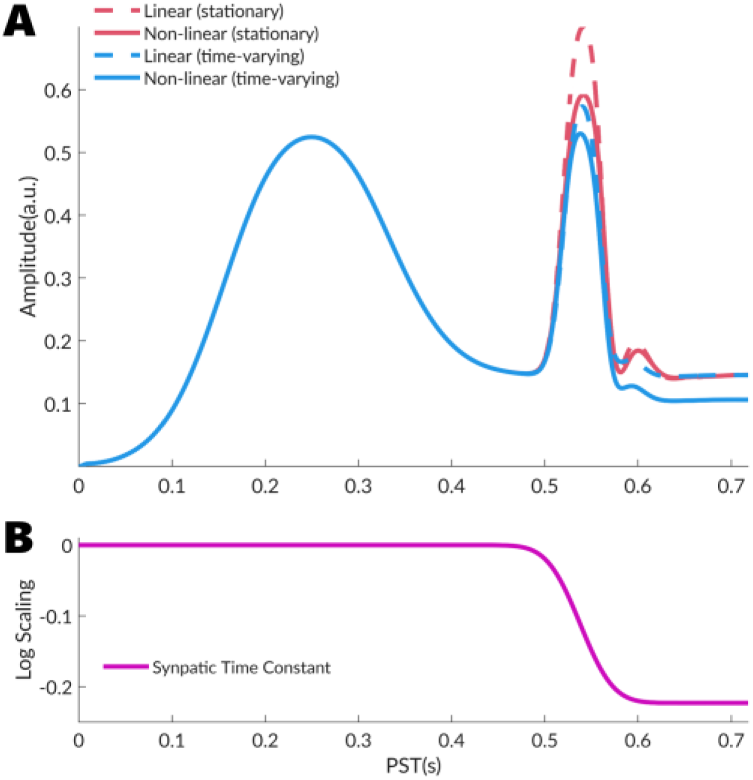
Temporal memory effects in sequential stimulus processing: Simulated LFP responses from a canonical microcircuit neural mass model to two sequential impulse stimuli separated by 300ms (onsets at 240ms and 540ms). Panel A contrasts linear predictions (dashed lines) obtained by summing individual stimulus responses against nonlinear dynamics (solid lines) from continuous integration across both events. Red traces show stationary parameter configurations where all synaptic time constants remain fixed throughout the epoch. Nonlinear dynamics (red solid) deviate from linear summation (red dashed) due to history-dependent carryover effects inherent in the neural mass equation. The system eventually returns to its original baseline, reflecting transient state perturbations within a fixed parameter regime. Blue traces incorporate time-varying parameters where synaptic time constants undergo a regime shift following the first stimulus. The nonlinear time-varying response (blue solid) shows enhanced deviation from linearity and settles to a persistently altered baseline rather than recovering to the initial state, demonstrating path-dependent hysteresis. The linear time-varying prediction (blue dashed) sums responses simulated independently with regime-appropriate static parameters (first regime for first stimulus, second regime for second stimulus), which returns to baseline because it fails to capture the continuous parameter trajectory. Panel B displays the evolution of synaptic time constants (log-scaled) for the time-varying case, revealing a sharp regime shift at approximately 500ms that persists throughout the remainder of the epoch, providing the mechanistic basis for the hysteretic baseline shift observed in Panel A.

###### Path-dependent hysteresis through parameter evolution

DCM-SR introduces a second, qualitatively distinct form of temporal memory operating on slower timescales and reflecting fundamentally different neurophysiological processes, including short-term synaptic plasticity, neuromodulation, and cognitive control processes reconfiguring connectivity patterns. When network parameters change following the first stimulus, the system responds to subsequent stimuli from an altered parameter regime, not merely from a transiently perturbed neural state. This is a form of hysteresis, where the system’s trajectory through parameter space determines which regime it occupies, and crucially, the system does not spontaneously return to its original configuration. Figure 2 (blue traces) illustrates this: with time-varying parameters, the response to sequential stimuli (solid line) shows greater deviation from linearity than the stationary case and settles to a persistently altered baseline rather than recovering to its initial state. The lower panel reveals the underlying mechanism: synaptic time constants decrease following the first stimulus and remain at this reduced value, reflecting a stable shift in the system’s dynamical landscape. In contrast, the linear summation (dashed line), computed by independently simulating each event with its respective static parameter regime, returns to baseline because it fails to capture the continuous parameter trajectory and its persistent effects.

This distinction between transient history dependence and persistent path dependence proves crucial for sequential paradigms where cognitive processes actively reconfigure network properties between events. By maintaining continuous neural states and permitting parameter evolution, DCM-SR naturally accommodates both forms of temporal memory, enabling disambiguation of whether differences in sequential responses arise from transient refractory effects or sustained changes in operating regime.

##### 2.1.1.3 Transition drivers

We now turn to the processes that induce transitions between parameter regimes. DCM-SR accommodates diverse inducers of network reconfiguration, coarsely distinguished into two classes. The first comprises predominantly exogenously-induced transitions time-locked to experimental events. The second comprises predominantly endogenously induced transitions arising from internal processes whose temporal profile is more ambiguous. Crucially, DCM-SR does not introduce fundamentally new mechanisms but extends the temporal expressiveness of established DCM operations. For instance, conventional DCM captures modulatory processes through B-matrix parameters encoding between-epoch effects. DCM-SR permits these same mechanisms to vary smoothly across time within a single epoch through time-varying A-matrix (and B-matrix) parameters, enabling characterisation of when and how network reconfigurations occur during sequential processing.

###### Exogenously induced transitions

External sensory perturbations provide the most straightforward class of transition inducers, where parameter changes are time-locked to experimentally controlled events. The nature of these transitions may reflect integrative processes that accumulate gradually across events or drastic reconfigurations that rapidly shift network dynamics. Cumulative parameter changes arise in tasks requiring incremental processing across sequential events. For instance, working memory maintenance is typically understood in terms of sustained activity within recurrent prefrontal and parietal networks (Olesen, Westerberg, and Klingberg 2004; Compte et al. 2000; Miller et al. 2003). Time-varying parameters could capture this through progressive lengthening of population time constants as items accumulate, or through evolving intrinsic connection strengths reflecting changes in recurrent stability. In contrast, discrete regime shifts occur when events trigger rapid network-wide reorganisations. Task switching exemplifies this, where instructional cues prompt abrupt reconfiguration of feedforward and feedback connectivity (Monsell 2003; Dove et al. 2000; Hyafil, Summerfield, and Koechlin 2009). By estimating set points within each parameter regime, DCM-SR can capture the large-scale shifts in effective connectivity and population dynamics that distinguish one task set from another. Figure 3 demonstrates simulated responses to sequential stimuli where transitions are aligned to event onsets, contrasting cumulative parameter changes (red trace, large *σ*_*i*_) versus discrete regime shifts (blue trace, small *σ*_*i*_).

**Figure 3.**
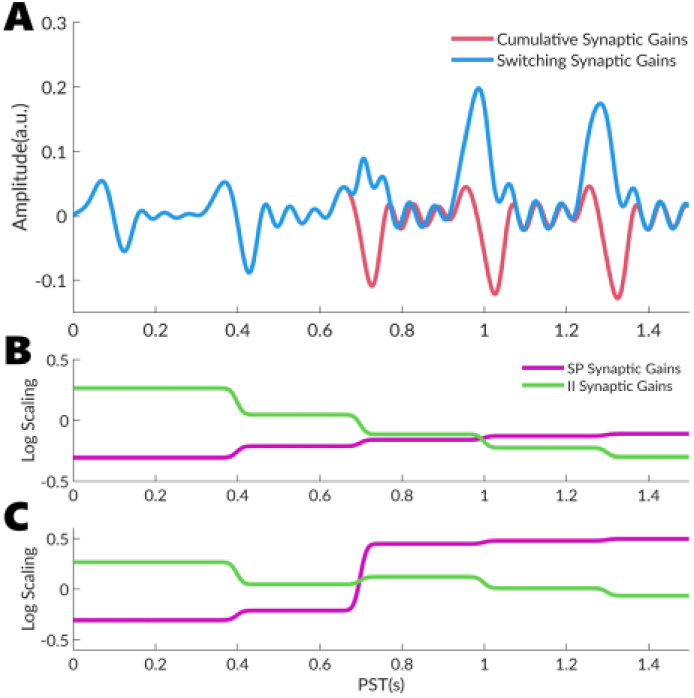
Exogenous transition types in DCM-SR. (A) Simulated responses to five identical sensory perturbations (blue markers) under cumulative (red) versus switching (blue) parameter regimes. (B) Cumulative exogenous transitions: Gradual, monotonic parameter evolution across repeated perturbations. Synaptic gains drift progressively with each stimulus presentation, consistent with processes such as working memory loading, repetition suppression, or learning. (C) Switching exogenous transitions: Abrupt parameter reconfiguration following the third perturbation. Synaptic gains undergo a discrete regime shift, with superficial pyramidal (SP) gains increasing and inhibitory interneuron (II) gains decreasing, consistent with the system crossing a stability boundary or undergoing a bifurcation. Both transition types are triggered by external sensory perturbations but exhibit qualitatively different temporal dynamics. Cumulative transitions reflect integrative processes that accumulate across events, whilst switching transitions reflect threshold-crossing or state-change mechanisms.

###### Endogenously induced transitions

Internal processes represent a more covert class of transition inducers where parameter reconfigurations arise from autonomous neural dynamics, exhibiting greater temporal uncertainty than exogenous transitions. This uncertainty is accommodated by the trajectory function parameterisation, which estimates both transition timings and dispersion. A canonical example is attentional deployment, which can occur either prospectively (Posner 1980; Chun 2000) or retrospectively (Thibault et al. 2016; Wildegger, Humphreys, and Nobre 2016). Conventional DCM cannot distinguish between these temporal profiles, as it captures attentional processes through stationary B-matrix modulation that distinguishes attended versus unattended conditions as a between-epoch contrast, conflating the temporal dynamics of when attention emerges. DCM-SR resolves this by estimating the progressive, time-varying engagement of attentional modulation within an epoch, permitting principled disambiguation between prospective and retrospective mechanisms. Figure 4 demonstrates this through simulated responses where endogenous parameter transitions occur at different times relative to stimulus onset, showing neural trajectories under four schemes: stationary parameters (black dashed), prospective time-variation where parameters change before the second stimulus (red), retrospective time-variation where parameters change after the first stimulus (blue), and combined dynamics (green dashed).

**Figure 4.**
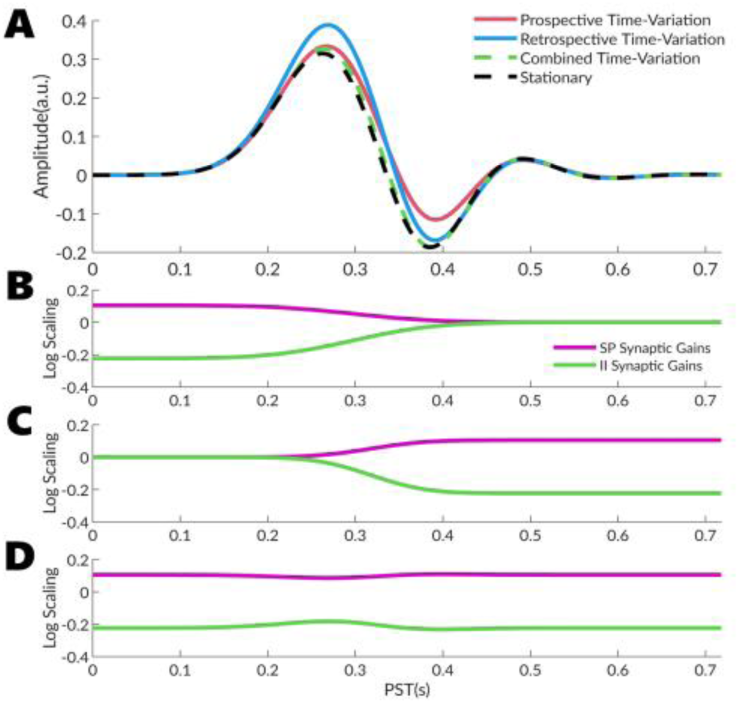
Endogenous transition types in DCM-SR. (A) Simulated responses to a single sensory perturbation at 300ms under prospective (red), retrospective (blue), combined (green dashed), and stationary (black dashed) parameter regimes. (B) Prospective endogenous transitions: Synaptic gain modulation occurs before the sensory perturbation, with superficial pyramidal (SP) gains decreasing and inhibitory interneuron (II) gains increasing. This preparatory reconfiguration enhances the early sensory-evoked response, consistent with anticipatory attention or predictive processing mechanisms. (C) Retrospective endogenous transitions: Synaptic gain modulation occurs after the sensory perturbation, with SP gains increasing and II gains decreasing. This sustained reconfiguration modulates the later components of the evoked response, consistent with working memory maintenance or retrospective attentional selection. (D) Combined prospective and retrospective transitions: When both transition types are present, parameters evolve continuously spanning the sensory event. In this simulation, the combined regime produces responses similar to the stationary condition, demonstrating that opposing gain changes (prospective decreases and retrospective increases) can yield net effects comparable to maintaining constant parameters.

###### Slow cortical potentials

Sequential paradigms frequently involve processes that bridge exogenous and endogenous characteristics, where brief external triggers initiate prolonged autonomous dynamics. Slow cortical potentials such as the contingent negative variation during motor preparation and the readiness potential preceding voluntary movement exemplify such mixed processes: an external cue or internal decision triggers a network state change, after which self-sustaining thalamocortical dynamics produce protracted accumulation spanning hundreds of milliseconds to seconds. These potentials reflect ramping activity maintained through recurrent connectivity, with ascending thalamic projections sustaining depolarisation of superficial pyramidal populations whilst reciprocal cortico-thalamic feedback stabilises the accumulated state. To model such dynamics, DCM-SR extends the repertoire of driving input functions beyond symmetric Gaussian bumps suited to brief transient stimuli. The skew-normal bump function enables asymmetric temporal profiles characteristic of anticipatory processes, combining a Gaussian probability density function *φ*(⋅) with a cumulative distribution function *ϕ*(⋅):

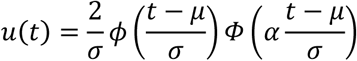

where *μ* represents the input peak time, *σ* controls temporal width, and *α* determines skewness. Negative *α* produces extended left tails (slow onset, rapid offset), positive *α* yields extended right tails (rapid onset, slow offset), whilst *α*=0 recovers the symmetric Gaussian. Figure 5 illustrates these asymmetric profiles across different *α* values and shows how combining skewed inputs with time-varying parameters produces slow cortical potential dynamics, where an initial brief input triggers parameter changes that enable sustained ramping activity culminating in a larger delayed response. This capacity to model mixed exogenous-endogenous processes proves essential for paradigms involving anticipatory preparation, decision accumulation, and sustained cognitive operations that begin with external triggers but unfold through intrinsic network dynamics.

**Figure 5.**
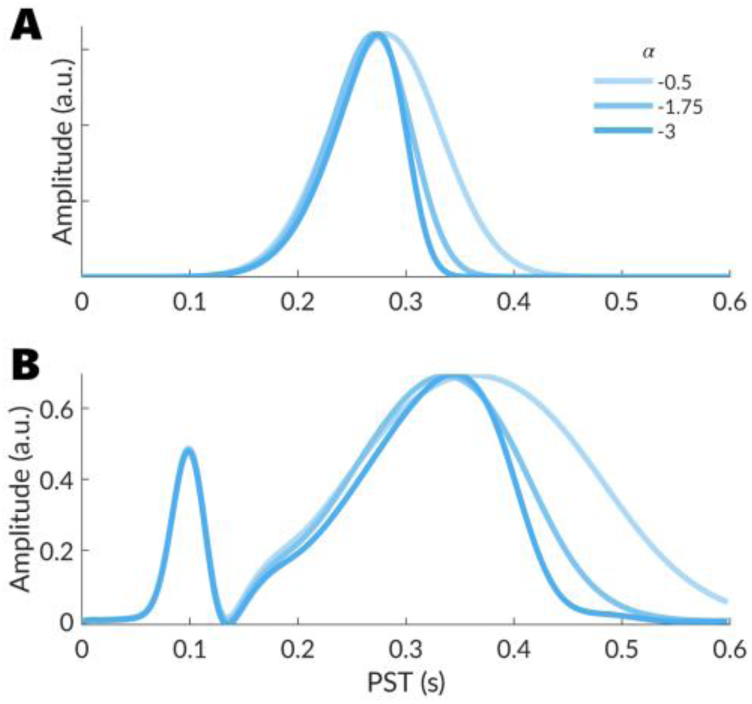
Slow cortical potential modelling with skewed Gaussian input function: (A) Skewed Gaussian input functions with varying asymmetry parameter *α*. As *α* becomes more negative (darker blue), the function exhibits greater positive skew, with a slower rise and sharper fall. (B) Simulated neural responses to two perturbations: a brief exogenous stimulus at 100ms and a slow, skewed input at 300ms with varying *α* values. The highly skewed input with *α* = −3 (darkest blue) produces a slow-ramping response characteristic of the contingent negative variation (CNV), with gradual amplitude build-up followed by sharp reduction. Less negative *α* values (lighter blue) produce more symmetrical responses. This parameterisation allows DCM-SR to capture preparatory slow cortical potentials observed in tasks requiring anticipation or temporal prediction.

### 2.2 Simulation Validity

To establish face validity for DCM-SR, we performed simulation studies using known ground truth to examine the consequences of substantial expansion in model dimensionality on model comparison and parameter identification. The parameter expansion creates competing risks: the enlarged parameter space may enable overfitting, spuriously favouring unnecessarily complex models, or conversely, the increased complexity may reduce sensitivity to genuine temporal dynamics, favouring inappropriate parsimony. Our simulations establish whether DCM-SR achieves accurate parameter recovery and correct model discrimination when parameters truly vary over time.

We conducted two complementary simulation studies. Simulation One examined whether model selection correctly discriminated between models differing with time-varying or stationary synaptic time constants. Simulation Two examined whether model selection correctly identified parameter transition drivers, disambiguating endogenous from exogenous perturbations. Together, these established that DCM-SR implements appropriate model selection, distinguishing intrinsic parameter variation from stimulus-evoked effects.

#### 2.2.1 General configuration

Both simulations employed a two-source network modelled with a canonical microcircuit. Neural activity was observed through two simulated sensors over 720 ms epochs sampled at 166 Hz. Synthetic observation noise was generated as an autoregressive process, AR(0.5), scaled to 20 dB signal-to-noise ratio. Ground-truth parameters were randomly sampled across 64 independent replications (Table 1). For each replication, we generated synthetic data from the ground-truth model and inverted both the true and alternative model specifications. Model selection performance was quantified through free energy differences, with Bayes factors greater than 3 indicating evidence favouring one model over another. We calculated the proportion of inversions correctly identifying the true generative model. Parameter recovery was assessed using standardised root mean square error (sRMSE) between estimated and ground-truth modulatory connectivity parameters, standardised by ground-truth standard deviation and averaged across replications.

**Table 1:** Sampling distributions for ground truth parameters in sequential DCM simulations.

| Parameter | Description | Sampling distribution |
| --- | --- | --- |
| A{1} | Extrinsic connection (forward) | $\mathcal{N}(1.5, 1/32)$ |
| A{2} | Extrinsic connection (forward) | $\mathcal{N}(1.2, 1/32)$ |
| A{3} | Extrinsic connection (backward) | $\mathcal{N}(1.5, 1/32)$ |
| A{4} | Extrinsic connection (backward) | $\mathcal{N}(1.2, 1/32)$ |
| G(1) | Intrinsic connection (SP gain) | $\mathcal{N}(0.9, 1/32)$ |
| G(2) | Intrinsic connection | $\mathcal{N}(1.0, 1/32)$ |
| G(3) | Intrinsic connection | $\mathcal{N}(1.1, 1/32)$ |
| G(4) | Intrinsic connection (II gain) | $\mathcal{N}(0.9, 1/32)$ |
| B(1,1) | Modulated connection (SP gain: source 1) | $\mathcal{N}(0.8, 1/8)$ |
| B(1,2) | Modulated connection (extrinsic forward: source 1) | $\mathcal{N}(0.3, 1/8)$ |
| B(2,1) | Modulated connection (extrinsic backward: source 1) | $\mathcal{N}(0.3, 1/8)$ |
| B(2,2) | Modulated connection (SP gain: source 2) | $\mathcal{N}(0.8, 1/8)$ |
| T(1) | Synaptic time constant (SS) | $\mathcal{N}(1, 1/32)$ |
| T(2) | Synaptic time constant (SP) | $\mathcal{N}(1, 1/32)$ |
| T(3) | Synaptic time constant (II) | $\mathcal{N}(1, 1/32)$ |
| T(4) | Synaptic time constant (DP) | $\mathcal{N}(1, 1/32)$ |

#### 2.2.2 Simulation One

Simulation One examined whether model selection correctly identified time-varying synaptic time constants across sequential events. Time constants were selected because alterations in synaptic time constants and efficacies both modulate response amplitude and timing, creating potential parameter confounds during model inversion. This confound provides a stringent test of model selection’s ability to balance model complexity against parsimony when multiple parameters can explain the same dynamics. All models included time-varying intrinsic connections (G), baseline connectivity (A), and modulatory connectivity (B), with the critical distinction concerning synaptic time constants (T). In Simulation 1A, the ground-truth model maintained stationary time constants, whilst the alternative permitted time-varying T parameters. In Simulation 1B, this configuration was reversed: the ground-truth included time-varying time constants (modulated by 12.5% to 25% across the epoch) whilst the alternative constrained them to remain fixed. Sequential driving inputs were delivered at irregular intervals (100, 175, 325, 550 ms) with 32 ms duration, demonstrating that the method accommodated arbitrary inter-stimulus timing.

#### 2.2.3 Simulation Two

Simulation Two examined whether model selection correctly distinguished exogenous driving inputs from endogenous state transitions. Sequential events were specified at 100, 200, 500, and 600 ms, comprising alternating exogenous and endogenous transitions. Exogenous perturbations occurred at 100 and 500 ms through driving inputs, whilst endogenous transitions followed 100 ms later through correlated parameter variations (r = 0.9) with the preceding exogenous event parameters. This correlation emulated realistic parameter drift following exogenous events, creating a stringent test by making the two mechanisms difficult to distinguish. Both models included time-varying synaptic gains (G), baseline connectivity (A), and modulatory connectivity (B), with the critical distinction concerning event specification. In Simulation 2A, the ground-truth model correctly specified all four events with their true generative mechanisms, whilst the alternative specified only the two exogenous inputs with time-varying parameters between events. In Simulation 2B, this configuration was reversed: the ground-truth comprised two exogenous events only, whilst the alternative allowed for endogenously induced parameter shifts 100 ms after each exogenous perturbation.

### 2.3 Empirical Validity

To establish construct validity, we applied DCM-SR to empirical data from an auditory go/no-go paradigm with extended cue-to-target intervals. This paradigm comprised five distinct phases: trial onset auditory cue, an anticipatory period, motor preparation preceding the expected target, presentation of the auditory target cue, and motor execution or inhibition. This structure provided diverse transition types spanning both exogenous and endogenous mechanisms, as well as shifts in cognitive context from auditory processing through anticipatory planning to motor control. We examined construct validity by addressing questions regarding the neural generators of the contingent negative variation during anticipatory and motor preparation phases, and by examining time-varying parameters during motor execution and inhibition, focusing on the extrinsic network involving inferior frontal gyrus and pre-supplementary motor area, as well as intrinsic pathways within the basal ganglia.

#### 2.3.1 Empirical Dataset

We analysed an open-access auditory go/no-go dataset (https://openneuro.org/datasets/ds003690) comprising 64-channel EEG from 75 participants (36 younger adults: 23 ± 3 years, 29 female; 39 older adults: 60 ± 5 years, 31 female) with normal or corrected vision and hearing and no neurological or psychiatric history. Each trial presented an anticipatory cue followed by either a go signal (250 ms, 1700 Hz tone, 80 trials/run) or a no-go signal (250 ms, 1300 Hz tone, 20 trials/run) at approximately 67 dB(A), requiring key presses for go trials and response withholding for no-go trials. Cue-to-target intervals were randomised using truncated exponential distributions (1.5 to 4.1 s) whilst post-stimulus intervals ranged 5.2 to 15.5 s. EEG was recorded at 500 Hz using a 64-channel International 10-20 system (reference CPz, ground FPz), with each participant completing two approximately 8-minute runs. We analysed correctly performed go and no-go trials only, discarding behavioural errors and artefact-contaminated epochs.

#### 2.3.2 Preprocessing

Continuous EEG was imported into SPM25 (Wellcome Centre for Human Neuroimaging, London) and band-pass filtered (4th-order Butterworth, 0.1 to 30 Hz) to attenuate drift and high-frequency noise whilst preserving ERP morphology. Data were downsampled to 200 Hz and epoched from −1000 to +6000 ms relative to trial onset without baseline correction to preserve temporal alignment. We excluded epochs containing behavioural errors and rejected epochs exhibiting |z| > 7 relative to within-channel distribution. Channels were re-referenced to the common average. We corrected for variable cue-to-target intervals using event-related warping. For each participant and condition, we specified two sequential events (cue, target) with onsets at recorded stimulus times, represented as Gaussian functions (σ = 50 ms). We did not apply response-time warping to align motor responses in go trials. Whilst such warping would reduce temporal jitter in go trials, it cannot be applied to no-go trials where no motor response occurs. Preserving response-time variability in both conditions maintains temporal equivalence when modelling motor execution versus motor inhibition, ensuring that differences in parameter estimates reflect genuine mechanistic distinctions rather than differential temporal alignment. Warping functions used 8th-order DST basis sets with Tukey tapering (taper ratio 0.1), estimated through 18 interpolation steps. Trial-averaged ERPs were computed using distance-weighted averaging, where trials were weighted by temporal deviation from the template. Subject-averaged ERPs were then grand-averaged across participants for DCM analysis.

#### 2.3.3 DCM specification

DCM-SR was fitted to grand-averaged ERPs spanning −500 to +4000 ms relative to trial onset. The network comprised six sources spanning auditory, motor, and prefrontal regions (Table 2). Bilateral auditory cortices (right and left superior temporal gyrus, rSTG and lSTG) received auditory inputs and were modelled using canonical microcircuit (CMC) neural mass models. Motor regions included left primary motor cortex (lM1, modelled with the motor microcircuit MMC) and left pre-supplementary motor area (lpSMA, CMC). Right inferior frontal gyrus (rIFG, CMC) represented prefrontal control, whilst basal ganglia-thalamic nucleus (BGT) was included as a silent subcortical source. The spatial forward model employed equivalent current dipoles.

**Table 2:**
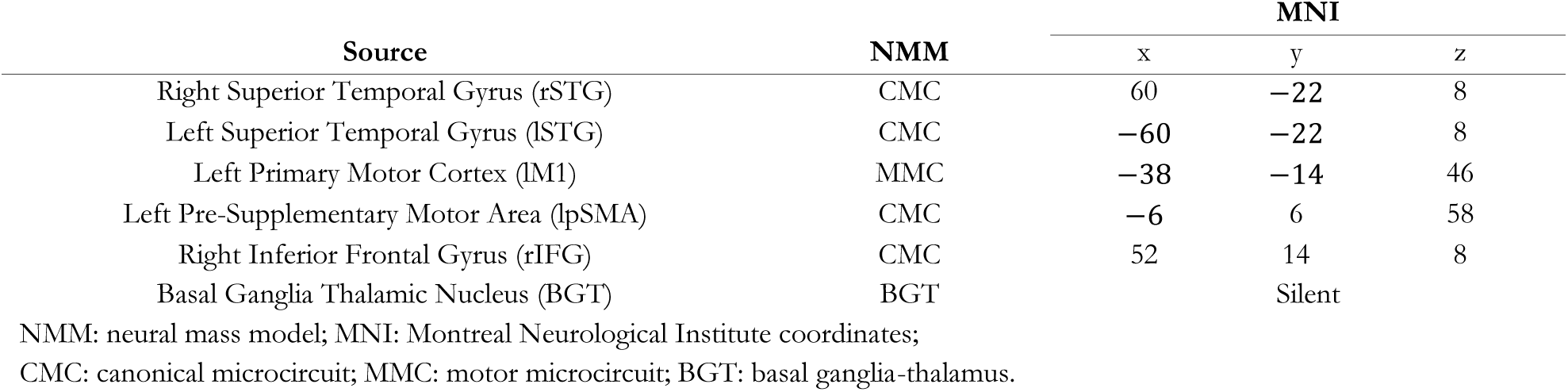
DCM-SR Source Specification.

The sequential structure specified five temporal events (Table 3), defined by where inputs are delivered and modulated connections (Figure 6). In addition to allowing extrinsic connection to be modulated by task effects, we permitted experimental modulation of the intrinsic connections within the BGT model to serve as proxy measures for the three primary functional pathways of the basal ganglia. Specifically, the model included modulatory parameters for the connection from the subthalamic nucleus to the internal globus pallidus (STN → GPi), representing the hyper-direct pathway; the striatum to the external globus pallidus (Striatum → GPe), representing the indirect pathway; and the striatum to the internal globus pallidus (Striatum → GPi), representing the direct pathway.

**Figure 6.**
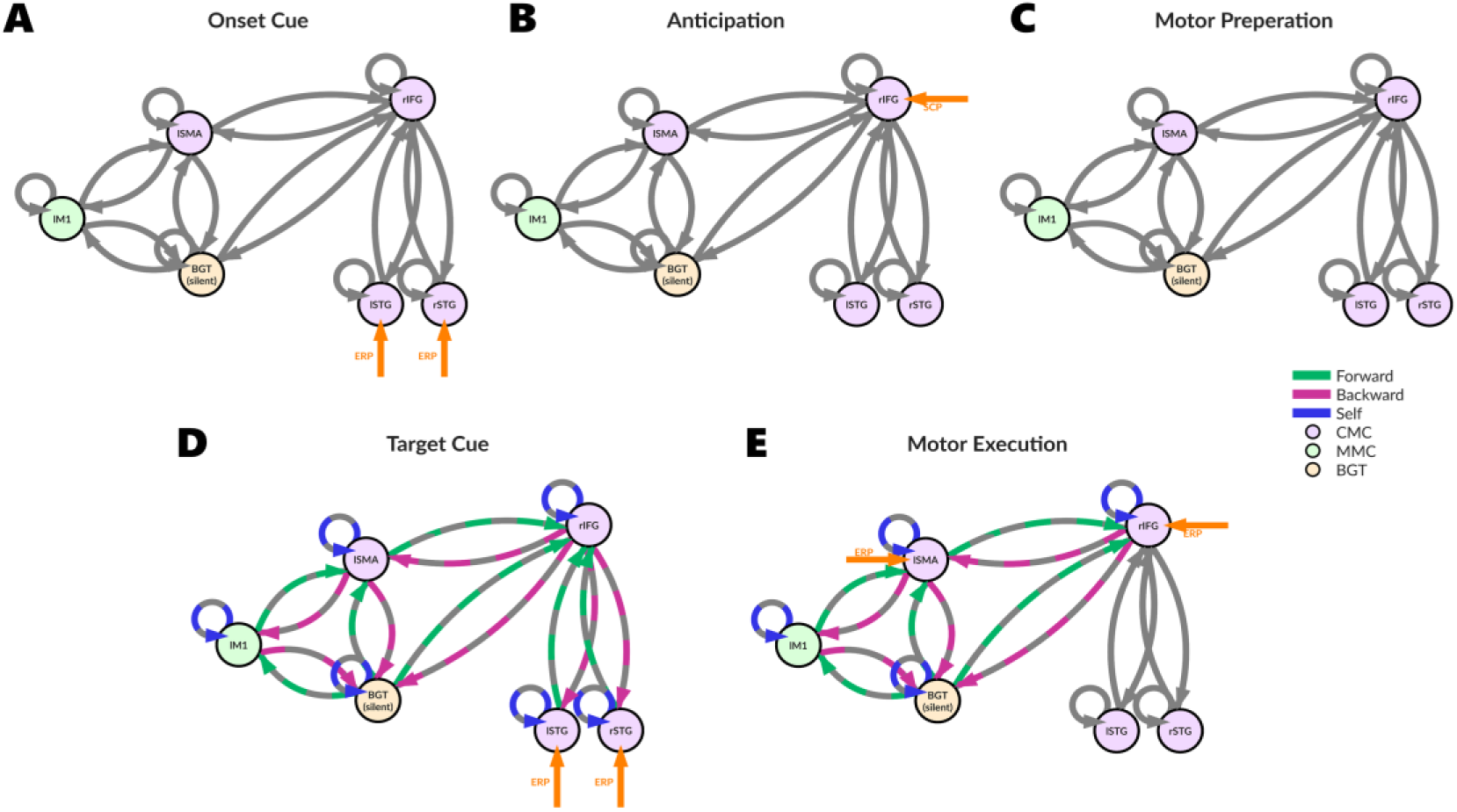
Network architecture and connectivity modulation across sequential events: Five panels (A–E) illustrate network configurations at onset cue (90 ms), anticipation (400 ms), motor preparation (1500 ms), target cue (1800 ms), and motor execution (2100 ms). Grey connections represent baseline extrinsic connectivity maintained throughout all events. Coloured connections (green: forward; magenta: backward; blue: self) indicate connections whose strengths are permitted to vary between go and no-go conditions. Orange arrows denote exogenous driving inputs modelled as bump functions for event-related potentials (ERP) and slow cortical potentials (SCP). Thick blue borders on nodes indicate regions with modulatory self-connections. Node types: CMC (canonical microcircuit), MMC (motor microcircuit), BGT (basal ganglia-thalamus). Abbreviations: rIFG, right inferior frontal gyrus; lpSMA, left pre-supplementary motor area; lM1, left primary motor cortex; BGT, basal ganglia-thalamus; lSTG, left superior temporal gyrus; rSTG, right superior temporal gyrus.

**Table 3:** DCM-SR Event Specification.

| Event | Onset (ms) | Duration (ms) | Bump Function | Input Node | Network Spec |
| --- | --- | --- | --- | --- | --- |
| Onset Cue | 90 | 16 | spm_erp_u | rSTG & lSTG | Fig 6A |
| Anticipation | 400 | 64 | spm_scp_u | rIFG | Fig 6B |
| Motor Preparation | 1500 | 32 | spm_null_u | Nil | Fig 6C |
| Target Cue | 1800 | 16 | spm_erp_u | rSTG & lSTG | Fig 6D |
| Motor Execution | 2100 | 24 | spm_erp_u | rIFG & lpSMA | Fig 6E |

All baseline extrinsic connectivity (A), modulatory connectivity (B), synaptic gains (G), and neural mass time constants (T) were permitted to vary across the five sequential states.

## 3 Results

In this section, we present validation of the DCM-SR framework through two complementary approaches. First, we establish face validity through controlled simulations that test the framework’s capacity for parameter recovery and model selection under known ground-truth conditions. Second, we demonstrate construct validity by applying the framework to empirical EEG data from a go/no-go task, examining whether recovered parameters reflect expected and plausible neural processes.

### 3.1 Simulation

The simulation study comprised four scenarios examining model identifiability. Simulation set one tested whether the framework correctly rejects unnecessary time-varying time constants (1A) and detects genuine temporal variation (1B). Simulation set two examined whether the framework detects endogenous synaptic changes (2A) and correctly rejects spurious endogenous transitions (2B). Model selection was quantified through free energy differences, with Bayes factors exceeding 3 indicating evidence favouring one model. Parameter recovery was assessed using sRMSE between estimated and ground-truth modulatory connectivity parameters across the full parameter trajectory, averaged across 64 replications.

#### 3.1.1 Simulation 1A

Simulation 1A tested whether the framework correctly rejects unnecessary time-varying time constants. The true generative model included time-varying G, A, and B parameters but fixed T parameters, whilst the alternative model allowed all parameters to vary with time.

Recovery of modulatory connectivity parameters was successful. Mean sRMSE for B parameters were: B(1,1) = 0.147 (95% CI [0.127, 0.167]), B(2,1) = 0.114 (95% CI [0.090, 0.137]), B(1,2) = 0.151 (95% CI [0.132, 0.170]), and B(2,2) = 0.288 (95% CI [0.246, 0.330]) (Figure 7A). Bayesian model comparison strongly favoured the true generative model, with an average free energy difference of 12.52 (95% CI [7.08, 17.95]) (Figure 7B). The correct model was identified in 84.4% of simulations (Figure 7 C). A representative simulation showed close correspondence between recovered and ground-truth parameters across all four sequential events (Figure 7D). Predicted responses accurately reconstructed both baseline and modulated LFP signals (Figure 8). The framework successfully distinguished the parsimonious model from the more complex alternative, correctly rejecting unnecessary temporal variation in T parameters when sequential dynamics were adequately explained by evolving synaptic efficacies alone.

**Figure 7.**
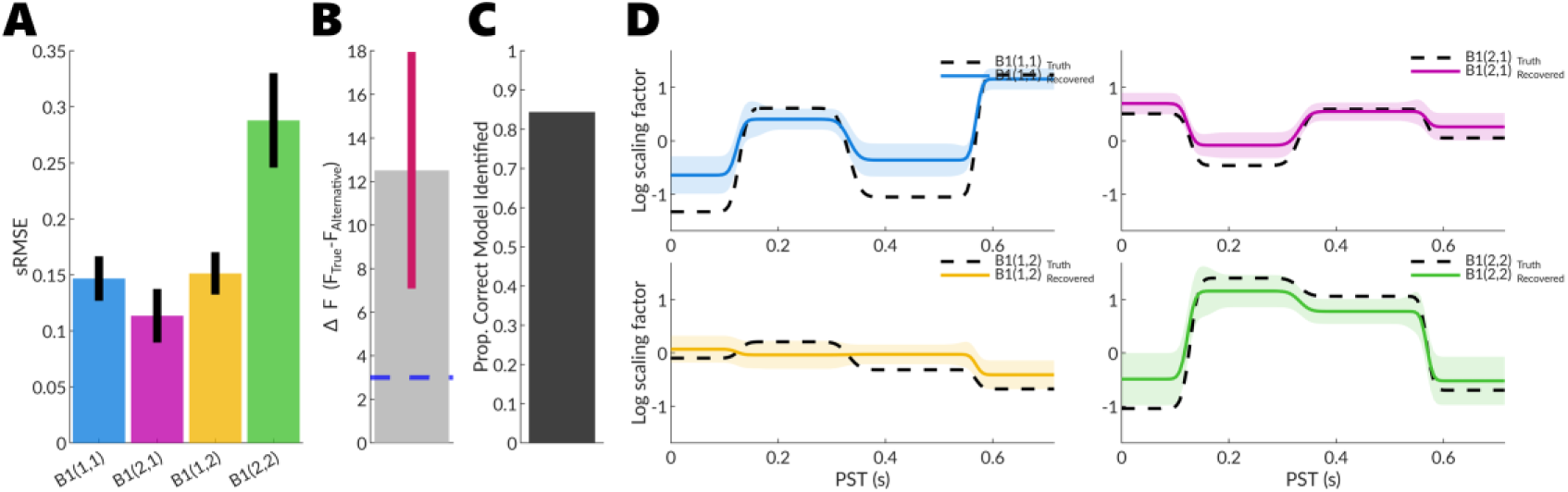
Parameter recovery analysis for Simulation 1A. A) Mean standardised root mean squared error (sRMSE) across simulations for recovered modulatory parameters B1(1,1), B1(2,1), B1(1,2), and B1(2,2), collapsed across time points. Error bars represent 95% confidence intervals. B) Model evidence comparison showing mean free energy (*ΔF*) across simulations. Grey bar represents the mean free energy difference between the true generative model (time-varying G, A, B parameters) and the alternative model (time-varying G, A, B, T parameters). Pink bar shows the 95% confidence interval. The blue horizontal line indicates the Bayes factor threshold of 3, demonstrating sufficient evidence to accept the true model. C) Model identification accuracy, showing the proportion of simulations in which the correct generative model was identified through Bayesian model comparison. D) Time-varying parameter estimates for modulatory connectivity parameters B1(1,1), B1(2,1), B1(1,2), and B1(2,2) from a representative simulation with sRMSE close to the mean. Dashed black lines represent ground truth values; coloured solid lines show recovered parameter estimates with shaded regions indicating posterior standard deviation. The B parameters represent experimentally modulated connectivity.

**Figure 8.**
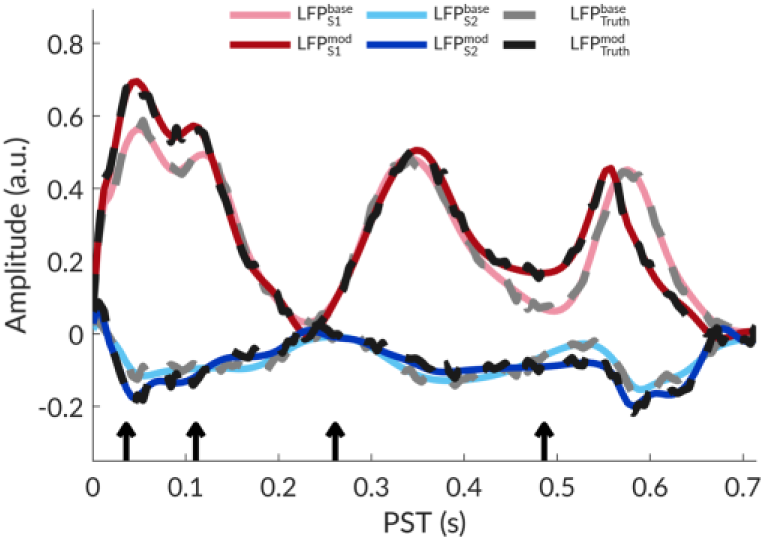
Representative simulated responses for Simulation 1A. Predicted local field potential (LFP) responses from two sources (S1 and S2) showing both baseline (base) and modulated (mod) activity. Grey and black lines represent the ground truth signals 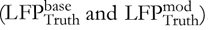. Coloured lines show the recovered responses from a representative simulation with root mean squared error close to the mean: pink and light blue lines represent baseline activity 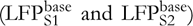, whilst dark red and dark blue lines represent modulated activity 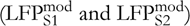. Black arrows indicate the timing of sequential stimulus events.

#### 3.1.2 Simulation 1B

Simulation 1B examined the framework’s ability to detect genuine temporal variation in synaptic time constants. The true generative model included time-varying G, A, B, and T parameters, with time constants modulated by ±16-22%, whilst the alternative model held T parameters fixed.

Recovery of modulatory connectivity parameters was successful. Mean sRMSE for B parameters were: B1(1,1) = 0.160 (95% CI [0.141, 0.179]), B1(2,1) = 0.118 (95% CI [0.100, 0.136]), B1(1,2) = 0.152 (95% CI [0.129, 0.174]), and B1(2,2) = 0.309 (95% CI [0.263, 0.354]) (Figure 9A), comparable to Simulation 1A. Bayesian model comparison favoured the true model, with a mean free energy difference of 7.71 (95% CI [5.88, 9.54]) (Figure 9B). The correct model was identified in 73.4% of simulations (Figure 9C), lower than Simulation 1A. This reduced performance likely reflects overlapping influences of time constants and synaptic efficacies on response dynamics. When time constants genuinely vary, the simpler alternative model remains competitive. Representative parameter estimates (Figure 9D) tracked ground-truth temporal structure with increased posterior uncertainty. Predicted responses (Figure 10) accurately reconstructed baseline and modulated LFP signals.

**Figure 9.**
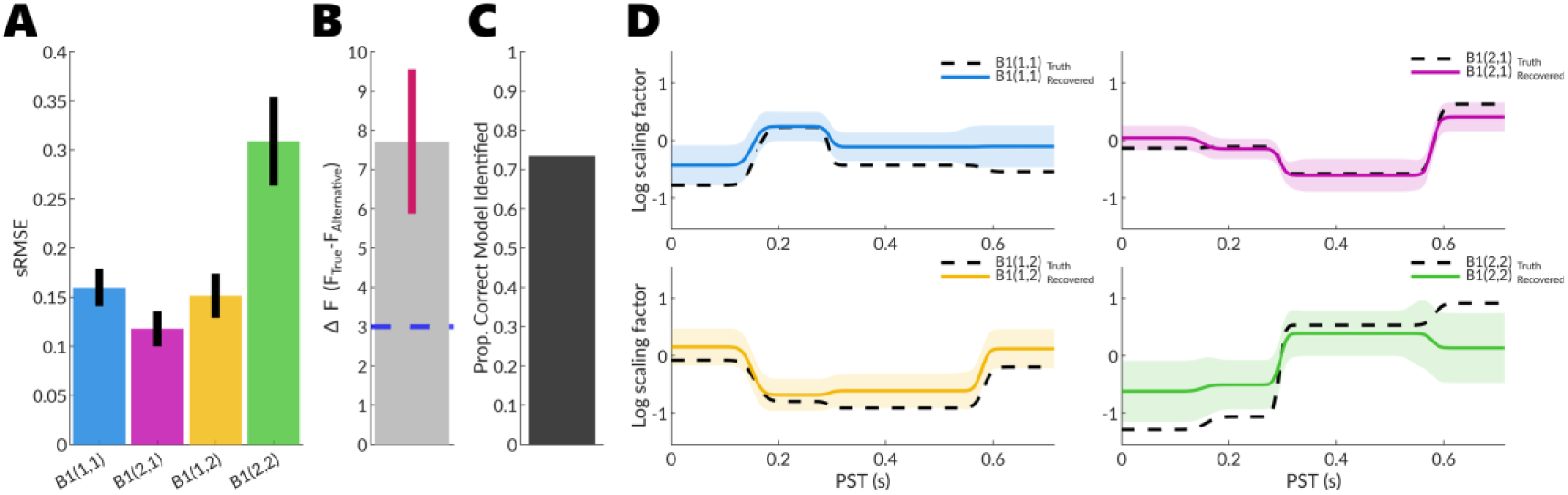
Parameter recovery analysis for Simulation 1B. A) Mean standardised root mean squared error (sRMSE) across simulations for recovered modulatory parameters B1(1,1), B1(2,1), B1(1,2), and B1(2,2), collapsed across time points. Error bars represent 95% confidence intervals. B) Model evidence comparison showing mean free energy (*ΔF*) across simulations. Grey bar represents the mean free energy difference between the true generative model (time-varying G, A, B, T parameters) and the alternative model (time-varying G, A, B parameters). Pink bar shows the 95% confidence interval. The blue horizontal line indicates the Bayes factor threshold of 3, demonstrating sufficient evidence to accept the true model. C) Model identification accuracy, showing the proportion of simulations in which the correct generative model was identified through Bayesian model comparison. D) Time-varying parameter estimates for modulatory connectivity parameters B1(1,1), B1(2,1), B1(1,2), and B1(2,2) from a representative simulation with sRMSE close to the mean. Dashed black lines represent ground truth values; coloured solid lines show recovered parameter estimates with shaded regions indicating posterior standard deviation. The B parameters represent experimentally modulated connectivity.

**Figure 10.**
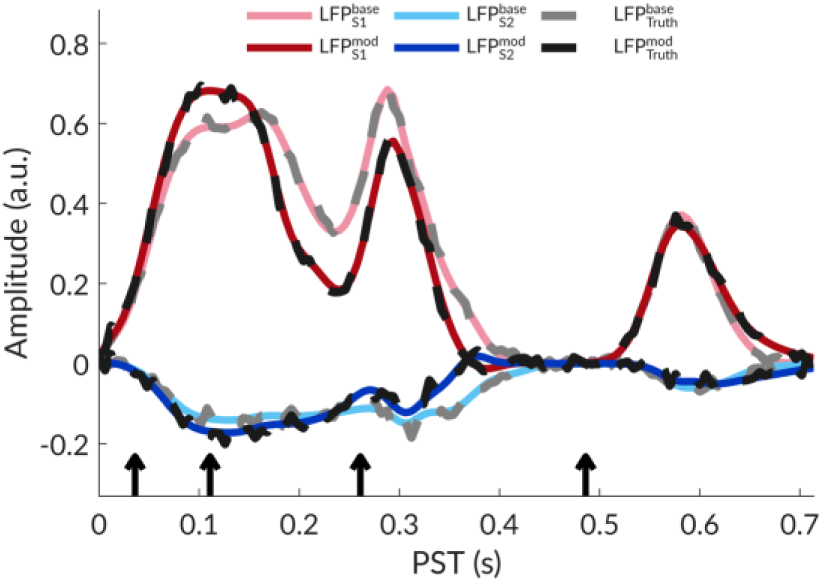
Representative simulated responses for Simulation 1B. Predicted local field potential (LFP) responses from two sources (S1 and S2) showing both baseline (base) and modulated (mod) activity. Grey and black lines represent the ground truth signals 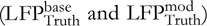. Coloured lines show the recovered responses from a representative simulation with root mean squared error close to the mean: pink and light blue lines represent baseline activity 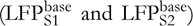, whilst dark red and dark blue lines represent modulated activity 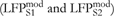. Black arrows indicate the timing of sequential stimulus events.

#### 3.1.3 Simulation 2A

Simulation 2A examined whether the framework detects endogenous synaptic changes. The true generative model specified four sequential events (100, 200, 500, 600 ms) with exogenous inputs at the first and third events. At the second and fourth events, parameters evolved endogenously with high autocorrelation (r = 0.9). The alternative model specified only two exogenous inputs (at 100 and 500 ms) with no endogenous evolution. Both models included time-varying G, A, and B parameters.

Recovery of modulatory connectivity parameters was successful. Mean sRMSE for B parameters were: B(1,1) = 0.261 (95% CI [0.221, 0.302]), B(2,1) = 0.244 (95% CI [0.189, 0.298]), B(1,2) = 0.213 (95% CI [0.167, 0.258]), and B(2,2) = 0.354 (95% CI [0.306, 0.402]) (Figure 11A). Bayesian model comparison favoured the true model, with a mean free energy difference of 23.15 (95% CI [12.63, 33.67]) (Figure 11B). However, the correct model was identified in 65.6% of simulations (Figure 11C), demonstrating only moderate sensitivity to endogenous changes. Representative parameter estimates (Figure 11D) tracked ground-truth values across all four events, including subtle endogenous transitions. Predicted responses (Figure 12) accurately reconstructed baseline and modulated LFP signals.

**Figure 11.**
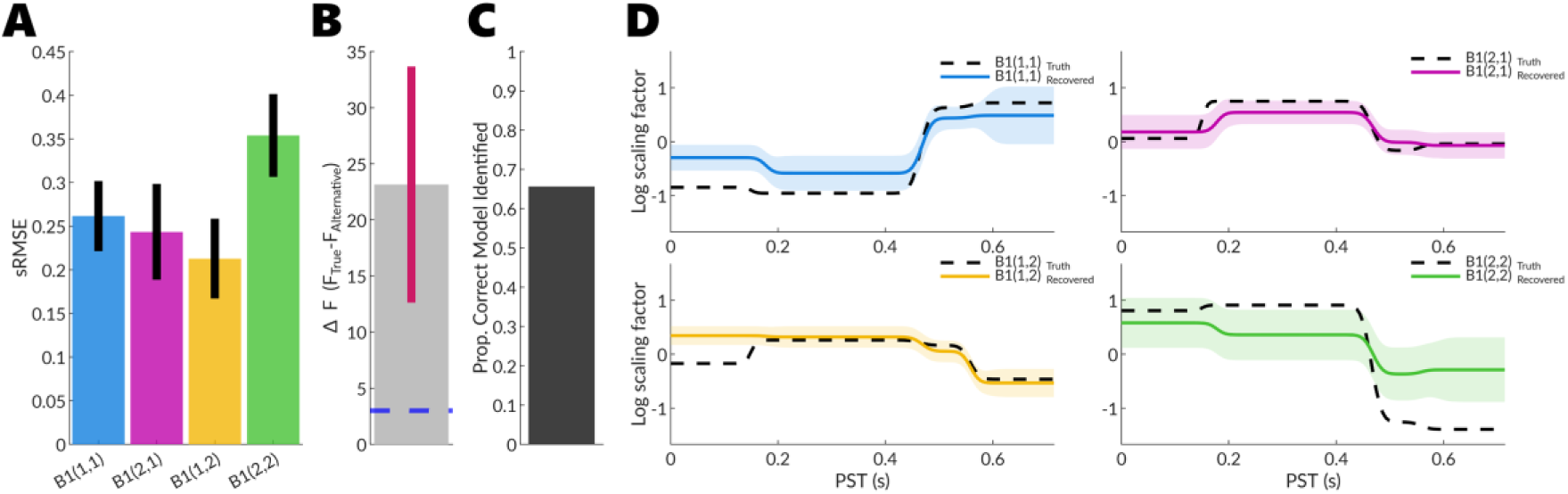
Parameter recovery analysis for Simulation 2A. A) Mean standardised root mean squared error (sRMSE) across simulations for recovered modulatory parameters B1(1,1), B1(2,1), B1(1,2), and B1(2,2), collapsed across time points. Error bars represent 95% confidence intervals. B) Model evidence comparison showing mean free energy (*ΔF*) across simulations. Grey bar represents the mean free energy difference between the true generative model (time-varying G, A, B, T parameters) and the alternative model (time-varying G, A, B parameters). Pink bar shows the 95% confidence interval. The blue horizontal line indicates the Bayes factor threshold of 3, demonstrating sufficient evidence to accept the true model. C) Model identification accuracy, showing the proportion of simulations in which the correct generative model was identified through Bayesian model comparison. D) Time-varying parameter estimates for modulatory connectivity parameters B1(1,1), B1(2,1), B1(1,2), and B1(2,2) from a representative simulation with sRMSE close to the mean. Dashed black lines represent ground truth values; coloured solid lines show recovered parameter estimates with shaded regions indicating posterior standard deviation. The B parameters represent experimentally modulated connectivity.

**Figure 12.**
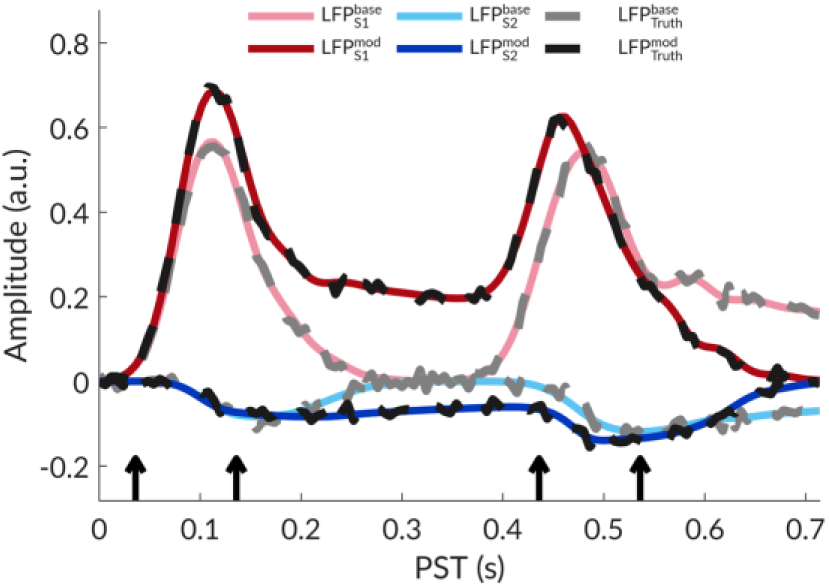
Representative simulated responses for Simulation 2A. Predicted local field potential (LFP) responses from two sources (S1 and S2) showing both baseline (base) and modulated (mod) activity. Grey and black lines represent the ground truth signals 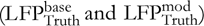. Coloured lines show the recovered responses from a representative simulation with root mean squared error close to the mean: pink and light blue lines represent baseline activity 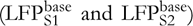, whilst dark red and dark blue lines represent modulated activity 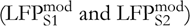. Black arrows indicate the timing of sequential stimulus events.

#### 3.1.4 Simulation 2B

Simulation 2B tested whether the framework correctly rejects spurious endogenous transitions when responses are entirely driven by exogenous inputs. The true generative model specified two sequential events (100 and 500 ms) with exogenous inputs at both time points. The alternative model specified four events (100, 200, 500, 600 ms) with exogenous inputs only at the first and third, incorrectly positing endogenous changes at 200 and 600 ms. Both models included time-varying G, A, and B parameters.

Recovery of modulatory connectivity parameters was successful. Mean sRMSE for B parameters were: B1(1,1) = 0.127 (95% CI [0.108, 0.146]), B1(2,1) = 0.078 (95% CI [0.065, 0.091]), B1(1,2) = 0.101 (95% CI [0.078, 0.125]), and B1(2,2) = 0.314 (95% CI [0.254, 0.374]) (Figure 13A), lower than Simulation 2A. Bayesian model comparison strongly favoured the true model, with a mean free energy difference of 11.41 (95% CI [10.29, 12.52]) (Figure 13B). The correct model was identified in 95.3% of simulations (Figure 13C), indicating that rejecting spurious endogenous changes is considerably easier than detecting genuine ones. Representative parameter estimates (Figure 13D) showed stable values between exogenous events. Predicted responses (Figure 14) accurately reconstructed LFP signals at both time points.

**Figure 13.**
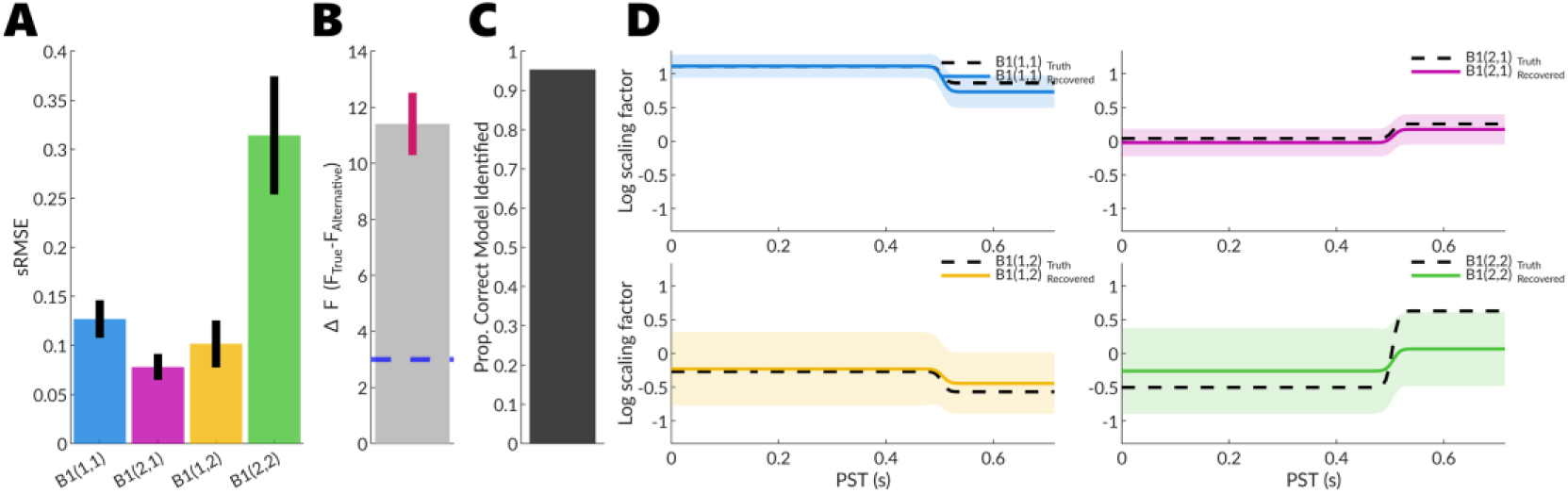
Parameter recovery analysis for Simulation 2B. A) Mean standardised root mean squared error (sRMSE) across simulations for recovered modulatory parameters B1(1,1), B1(2,1), B1(1,2), and B1(2,2), collapsed across time points. Error bars represent 95% confidence intervals. B) Model evidence comparison showing mean free energy (*ΔF*) across simulations. Grey bar represents the mean free energy difference between the true generative model (time-varying G, A, B, T parameters) and the alternative model (time-varying G, A, B parameters). Pink bar shows the 95% confidence interval. The blue horizontal line indicates the Bayes factor threshold of 3, demonstrating sufficient evidence to accept the true model. C) Model identification accuracy, showing the proportion of simulations in which the correct generative model was identified through Bayesian model comparison. D) Time-varying parameter estimates for modulatory connectivity parameters B1(1,1), B1(2,1), B1(1,2), and B1(2,2) from a representative simulation with sRMSE close to the mean. Dashed black lines represent ground truth values; coloured solid lines show recovered parameter estimates with shaded regions indicating posterior standard deviation. The B parameters represent experimentally modulated connectivity.

**Figure 14.**
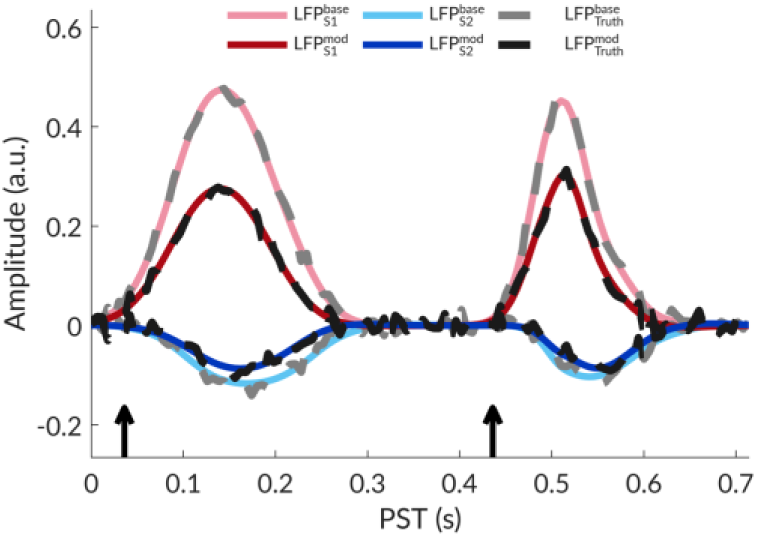
Representative simulated responses for Simulation 2B. Predicted local field potential (LFP) responses from two sources (S1 and S2) showing both baseline (base) and modulated (mod) activity. Grey and black lines represent the ground truth signals 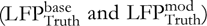. Coloured lines show the recovered responses from a representative simulation with root mean squared error close to the mean: pink and light blue lines represent baseline activity 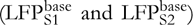, whilst dark red and dark blue lines represent modulated activity 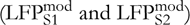. Black arrows indicate the timing of sequential stimulus events.

#### 3.1.5 Summary

These simulations establish face validity for DCM-SR, demonstrating robust parameter recovery and appropriate model selection. Parameter estimation remained accurate across all scenarios (mean sRMSE < 0.36). The framework exhibited conservatism, favouring parsimony unless data provided compelling evidence for complexity. Rejecting spurious complexity proved more reliable than detecting genuine complexity: spurious time-varying time constants were rejected in 84.4% of cases (Simulation 1A) versus 73.4% detection of genuine variation (Simulation 1B), whilst spurious endogenous transitions were rejected in 95.3% of cases (Simulation 2B) versus 65.6% detection of genuine endogenous changes (Simulation 2A). This asymmetry reflects the model comparison preference for simpler explanations, particularly when time constants and synaptic efficacies exert overlapping influences or when endogenous changes are subtle.

### 3.2 Empirical

Having established face validity through controlled simulations, we now apply the DCM-SR framework to EEG recordings from a go/no-go task. The go/no-go paradigm provides an ideal testbed because it engages well-characterised cortical and subcortical circuits across an extended temporal sequence. Following an orienting cue, participants maintain an attentive preparatory state during which the contingent negative variation (CNV) emerges as a slow cortical potential reflecting sustained anticipation. Upon target presentation, the task demands rapid response selection and, for no-go trials, suppression of prepotent motor responses through distributed fronto-basal ganglia networks. By tracking both fast-evolving neural states and slow-evolving connectivity parameters, we assess whether the framework captures established neurophysiological processes.

#### 3.2.1 Model fit to temporal modes

Following standard DCM procedures, the 64-channel data were projected onto eight principal modes via singular value decomposition. Figure 15 displays the model fit to the first three modes, comparing observed responses (dashed lines) against predicted responses (solid lines) for both conditions. Quantitative assessment across all eight modes yielded *R*^2^=91.44% for the go condition and *R*^2^=89.12% for the no-go condition, indicating that DCM-SR accounted for over 90% of variance in the observed temporal dynamics.

**Figure 15.**
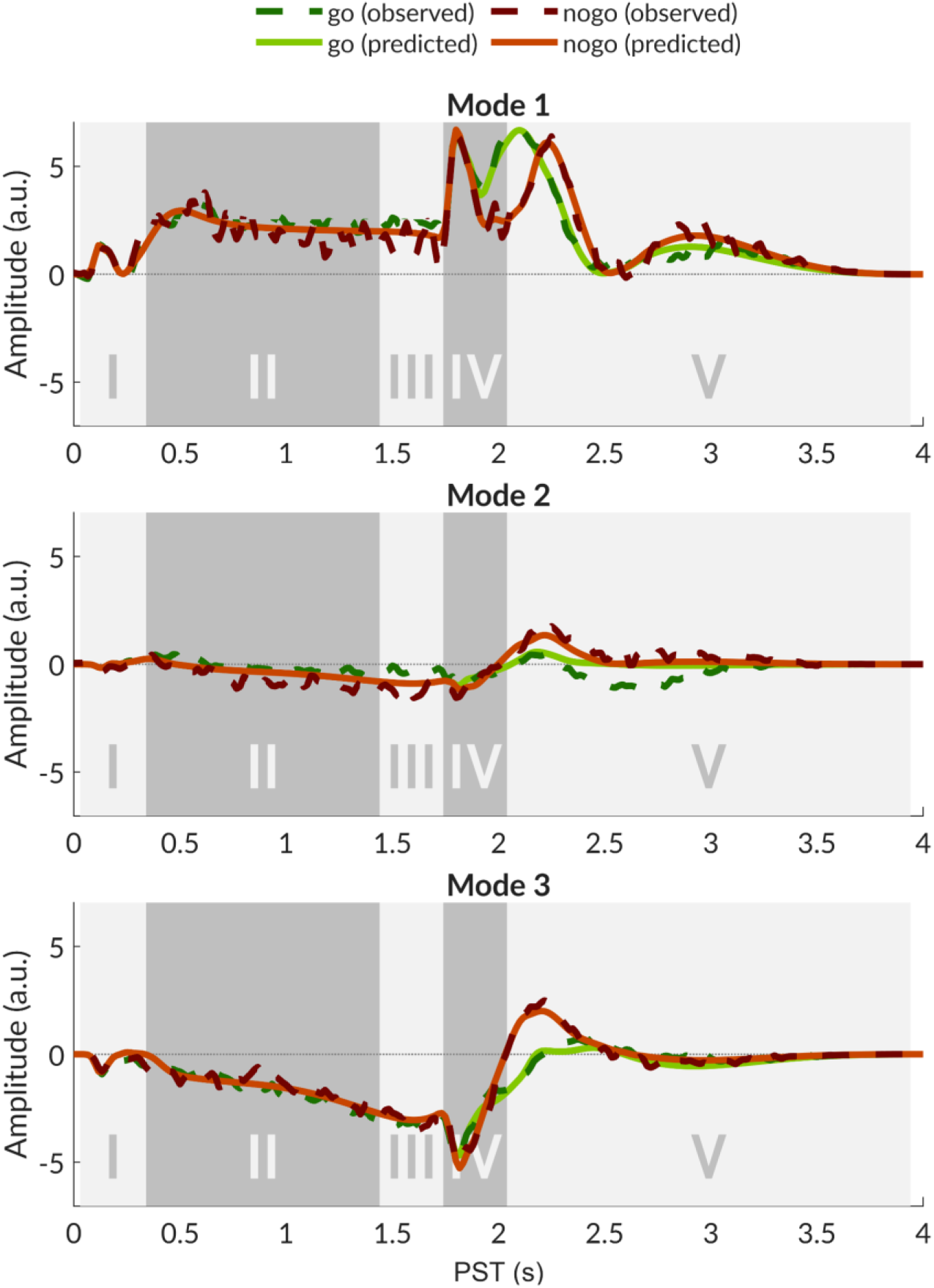
Model fit to temporal evolution of principal modes. Observed (dashed lines) and predicted (solid lines) temporal responses for the first three modes extracted via singular value decomposition of grand-averaged EEG data, shown separately for go (green) and no-go (red) conditions. Grey shaded regions delineate the five sequential experimental phases: (I) onset cue (90 ms), (II) anticipation (400--1800 ms), (III) motor preparation (1800 ms), (IV) target cue (1800 ms), and (V) motor execution (2100 ms onwards). DCM-SR achieved close correspondence between predicted and observed temporal dynamics across all phases, with coefficient of determination values of *R*^2^ = 91.44 (go condition) and *R*^2^ = 89.512 (no-go condition) computed across all eight temporal modes.

#### 3.2.2 Parameter and neural state trajectories

Having established that DCM-SR accurately captures observed electrophysiological responses, we examined the estimated parameters and neural state trajectories to establish construct validity. We focused on two phenomena: the neural generators of the CNV and motor response inhibition. The CNV is a slow negative potential arising during anticipatory preparation, extensively characterised in terms of its associations with cognitive processes and behavioural outcomes. However, mechanistic accounts linking these phenomena to underlying neural generators remain largely unexplored. Here, we demonstrate how DCM-SR can decompose this data feature into constituent neurophysiological processes. Conversely, motor response inhibition represents a well-studied phenomenon with established mechanisms, allowing us to assess whether DCM-SR generates physiologically plausible explanations. The following descriptions serve primarily to illustrate the explanatory accounts DCM-SR can provide, rather than constituting formal model-based evidence for specific mechanistic hypotheses. Nevertheless, examining whether recovered dynamics align with theoretical predictions provides essential construct validity.

##### 3.2.2.1 Contingent Negative Variation Generation

The CNV develops between a warning stimulus and an imperative stimulus requiring a behavioural response (Walter et al. 1964). The CNV comprises two functionally dissociable components: an early orienting phase reflecting attention allocation and arousal, maximal over frontal regions, and a late expectancy phase reflecting motor preparation, maximal over central regions, which shares substantial overlap with the readiness potential (Loveless and Sanford 1974; Gaillard 1977). Despite over 60 years since the CNV’s initial description, the neuronal generators of slow cortical potentials remain incompletely understood. The prevailing view holds that depolarisation from excitatory postsynaptic potentials at superficial-layer apical dendrites is the primary generator of slow negative potentials (Birbaumer et al. 1990). However, these proposed neural generators have been largely extrapolated from visually evoked current source density analyses in animal preparations (Mitzdorf 1985) and studies of slow potentials emerging from pathophysiological states such as seizures (Caspers, Speckmann, and Lehmenkühler 1980), rather than from direct recordings during cognitive tasks. More recently, theoretical work has suggested that synchronised afterhyperpolarisations in deep pyramidal cells may be the principal generator of slow potentials (Buzsáki, Anastassiou, and Koch 2012). The CNV, therefore, presents an opportunity to adjudicate between competing mechanistic accounts using recordings obtained during actual cognitive processing.

###### CNV Orienting and Early Cortical Dynamics (Early-to-Mid Phase II)

The earliest alterations following cue onset were characterised by rapid shifts in cortical neural states, primarily localised to the rIFG. As the primary cortical target for thalamo-cortical input, the rIFG exhibited early transient depolarisation across spiny stellate, inhibitory interneurone, and deep pyramidal populations (Figure 16). This initial activation occurred alongside broader suppression of the motor system, marked by hyperpolarisation of deep pyramidal (DP) populations in both the rIFG and the lpSMA. During this period, lM1 showed consistent hyperpolarising trends across all populations, most prominently within middle pyramidal (MP) and inhibitory interneurons (II) cells. Whilst these cortical shifts were pronounced, the basal ganglia-thalamic (BGT) populations maintained a relatively steady state near baseline.

**Figure 16.**
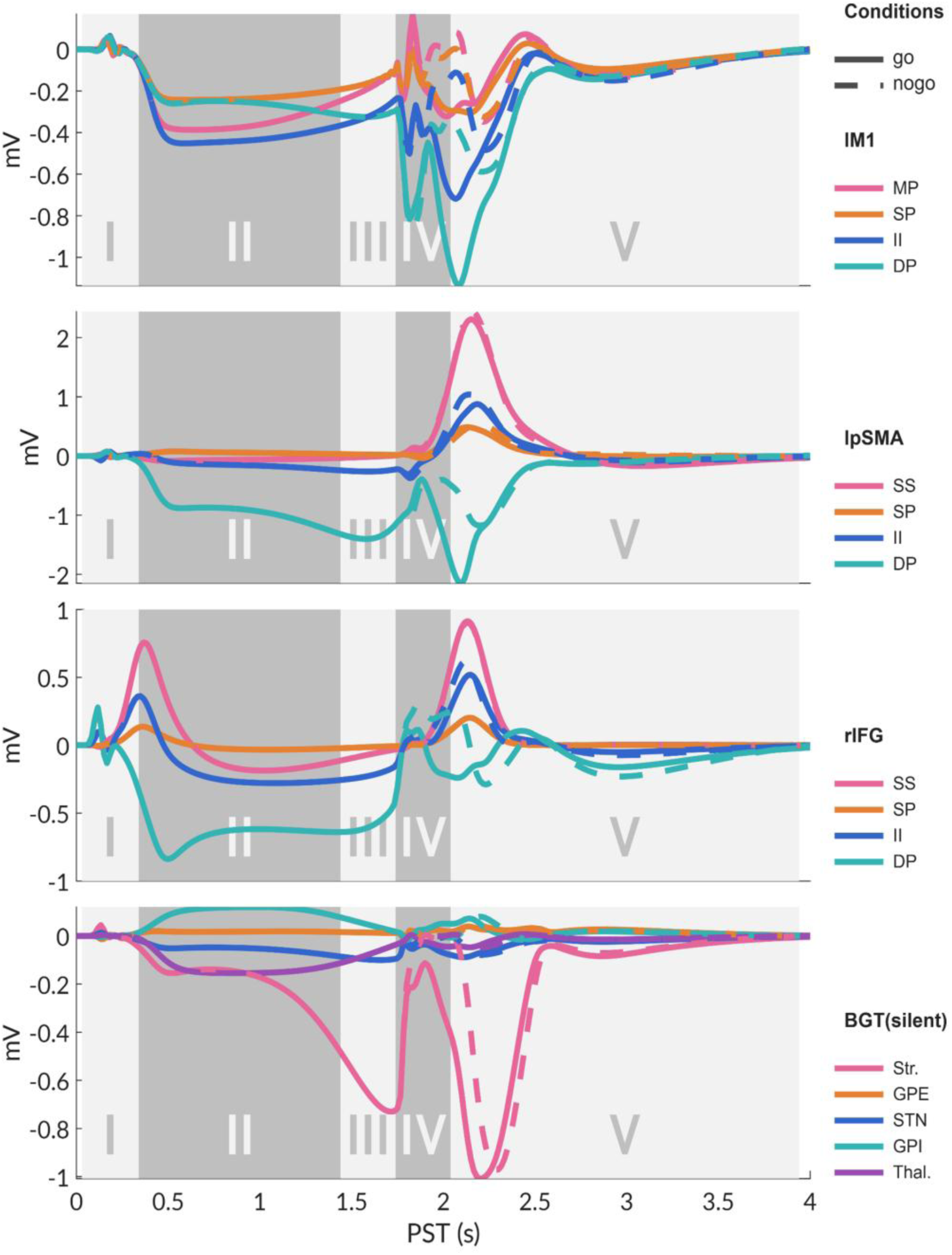
Temporal evolution of neural states across experimental phases. Posterior estimates of time-varying neural activity (depolarization in mV) across the fronto-striatal-thalamic network, illustrating the continuous evolution of neural states across five discrete experimental phases: (I) onset cue (90 ms), (II) anticipation (400--1800 ms), (III) motor preparation (1500 ms), (IV) target cue (1800 ms), and (V) motor execution (2100 ms onwards). Panels display conditional estimates for key network nodes: lM1 (left primary motor cortex), lpSMA (left pre-supplementary motor area), rIFG (right inferior frontal gyrus), and the BGT (basal ganglia-thalamus) system. Solid lines represent posterior expectations for the go condition and dashed lines represent the no-go condition. The BGT components are modeled as ‘silent’ neural populations, representing hidden states that contribute to circuit dynamics without directly mapping onto observed scalp data. Line colours distinguish specific neural populations: lM1, lpSMA, and rIFG: MP, middle pyramidal; SP, superficial pyramidal; II, inhibitory interneurons; DP, deep pyramidal; SS, spiny stellate. BGT: Str., striatum; GPe, globus pallidus externa; STN, subthalamic nucleus; GPi, globus pallidus interna; Thal., thalamus. Roman numerals (I--V) and shaded backgrounds denote the temporal boundaries of each experimental phase.

These evolving neural states were underpinned by specific modulations in time-varying extrinsic connectivity (Figure 17). The most prominent driver of early cortical activity was increased connection strength from the thalamic population of the BGT to the spiny stellate (SS) cells of the rIFG. This thalamo-cortical projection exhibited a rapid rise following the onset cue and reached a stable plateau maintained throughout the anticipatory period (Phase II). Within the internal circuitry of the basal ganglia (Figure 18), this inhibitory posture was further reinforced by a pronounced increase in hyperdirect pathway strength, specifically from the subthalamic nucleus (STN) to the globus pallidus interna (GPi). This rise in hyperdirect signalling occurred concurrently with a modest reduction in direct pathway activity from the striatum to the GPi, effects that collectively limit motor facilitation during the attentional orienting phase. In contrast, the indirect pathway via the globus pallidus externa (GPe) remained stable, suggesting that initial global motor suppression is primarily mediated by the hyperdirect pathway.

**Figure 17.**
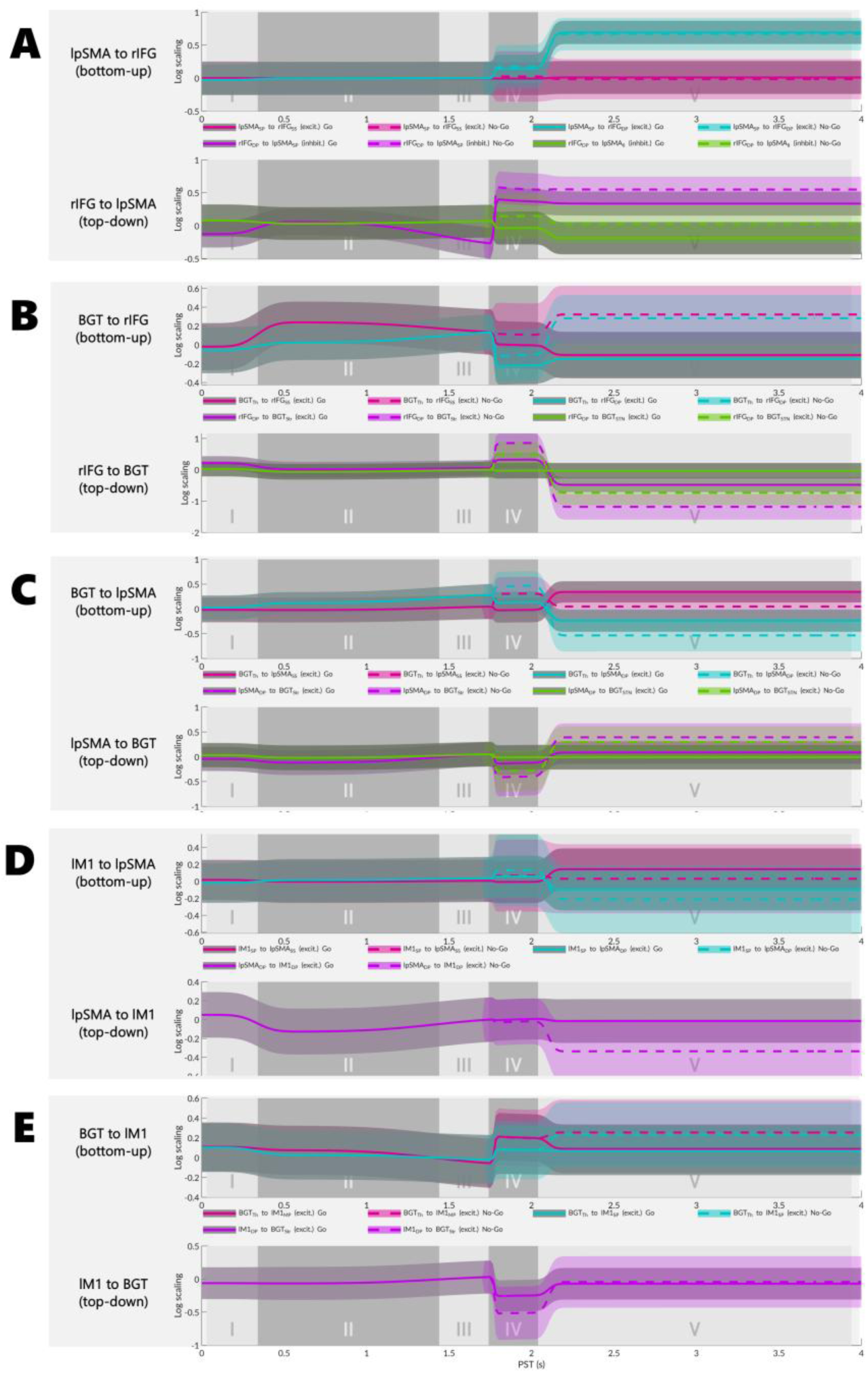
A. Network connectivity during target cue and motor execution phases: Posterior estimates for modulatory effects on extrinsic connectivity and self-connections at target cue (left) and motor execution (right) phases, showing connections permitted to differ between go and no-go conditions. These values reflected modulations from baseline connectivity (B matrix parameters). For each connection, the upper value indicates the posterior expectation (log-scaled modulation strength) and the lower value in parentheses indicates the posterior probability. Connections are colour-coded by type: forward (green), backward (magenta), and self-connections (blue). Node types denote canonical microcircuit (CMC), motor microcircuit (MMC), and basal ganglia-thalamus (BGT). Orange arrows indicate exogenous driving inputs (ERP). Abbreviations: rIFG, right inferior frontal gyrus; lSMA, left supplementary motor area; lM1, left primary motor cortex; BGT, basal ganglia-thalamus; lSTG, left superior temporal gyrus; rSTG, right superior temporal gyrus. B--D. Temporal evolution of extrinsic connectivity across experimental phases: Time-varying parameter trajectories showing modulatory effects on extrinsic connections between key network nodes across the five experimental phases: (I) onset cue (90 ms), (II) anticipation (400--1800 ms), (III) motor preparation (1500 ms), (IV) target cue (1800 ms), and (V) motor execution (2100 ms onwards). Neural population abbreviations: SS, spiny stellate; TH, thalamus; DP, deep pyramidal; STR, striatum; STN, subthalamic nucleus; SP, superficial pyramidal; II, inhibitory interneurons.

**Figure 18.**
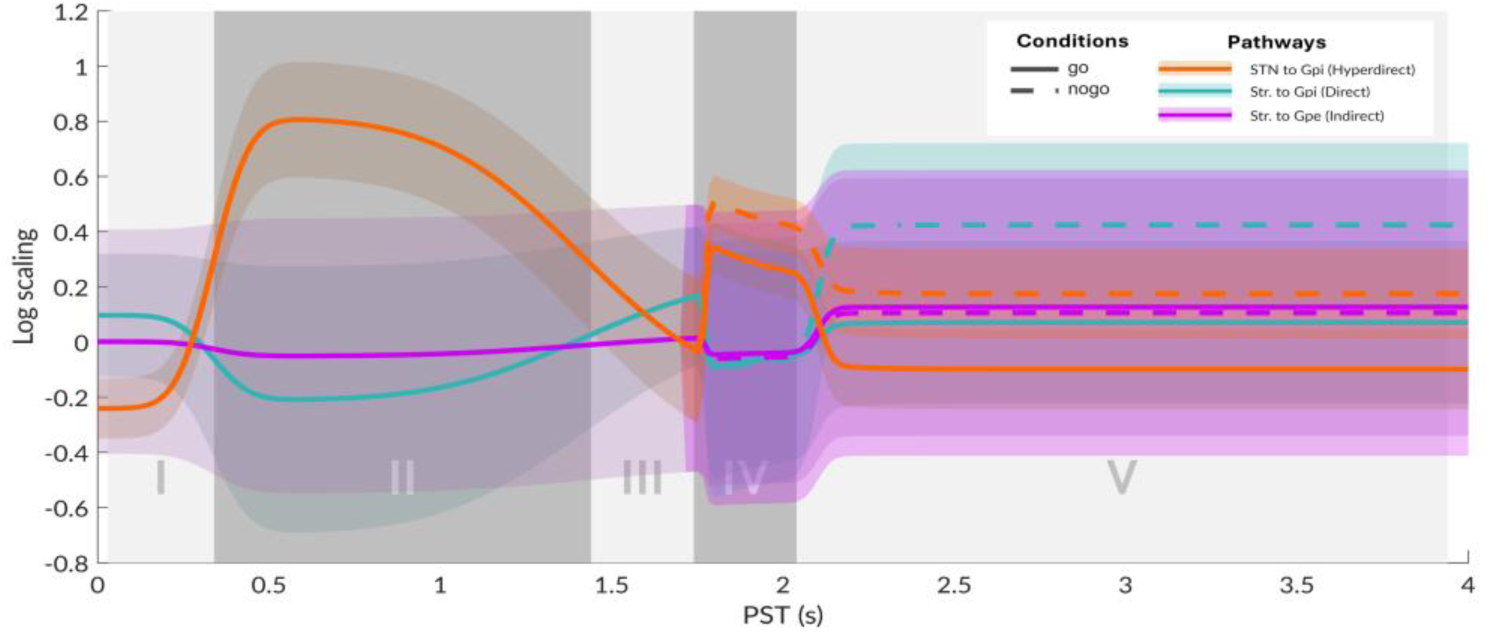
Temporal evolution of key basal ganglia intrinsic connections across experimental phases: Time-varying parameter trajectories show log-scaled modulations for the hyperdirect (STN to GPi; orange), direct (Str to GPi; teal), and indirect (Str to GPe; magenta) pathways. Solid lines represent posterior expectations for the go condition and dashed lines represent the nogo condition. Shaded regions indicate posterior standard deviations. The vertical panels represent the five experimental phases: (I) onset cue, (II) anticipation, (III) motor preparation, (IV) target cue, and (V) motor execution.

Whilst our modelling framework cannot definitively resolve the laminar source of the CNV, the observed hyperpolarisation of DP cells offers compelling evidence for accounts attributing slow cortical potentials to synchronised after-hyperpolarisations within DP populations (Buzsáki, Anastassiou, and Koch 2012), contrasting with classical emphasis on superficial-layer depolarisation (Birbaumer et al. 1990). The dominant connectivity change of sustained elevation of thalamo-cortical drive targeting superficial layers of prefrontal cortex is consistent with intracerebral recordings showing CNV-like activity in subcortical structures, including the putamen (Bareš and Rektor 2001) and evidence for basal ganglia-thalamo-cortical circuit involvement in CNV generation (Rektor et al. 2004). Together, these findings suggest basal ganglia circuits simultaneously enhance arousal in prefrontal regions via thalamic input to rIFG superficial layers whilst suppressing premature motor output via the hyperdirect pathway, with the surface-negative CNV potentially reflecting synchronised hyperpolarisation of DP populations.

###### CNV Expectancy (Late Phase II & Phase III)

As cortical processing transitioned from orienting to expectancy, the most prominent shifts occurred within the subcortical populations of the BGT system (Figure 16). The striatum exhibited marked hyperpolarisation, accompanied by two critical modulations in basal ganglia connectivity. First, the hyperdirect pathway (STN to GPi) showed substantial reduction from its Phase II peak, aligning with release of rapid global motor suppression typically engaged during early stimulus processing (Nambu, Tokuno, and Takada 2002). Concurrently, the direct pathway (Str to GPi) exhibited modest strengthening, representing enhanced striatal inhibition of the GPi and consequent disinhibition of thalamo-cortical circuits (Jonathan W. Mink 2003). Whilst the indirect pathway (Str to GPe) remained relatively stable, its influence on motor output was inherently constrained by the omission of the GPe-to-GPi projection within the current model. Collectively, these subcortical dynamics reflect progressive release of tonic motor inhibition designed to permit execution of prepared motor programmes (Jonathan W. Mink 1996).

Concomitant shifts across the cortical motor hierarchy revealed distinct laminar dissociation within lM1. In this region, increased depolarisation in the MP and SP populations occurred alongside heightened hyperpolarisation in the deep pyramidal cells and modest increases in inhibitory interneurone gain. This pattern suggests that active motor preparation in superficial layers is maintained concurrently with suppression of corticospinal output channels in deep layers until movement execution. The lpSMA exhibited slight deepening in deep pyramidal hyperpolarisation, whilst the rIFG maintained a relatively steady state. Notably, the most prominent cortico-cortical change was sharp reduction in connection strength from the rIFG to the lpSMA during late Phase III (Figure 17). This modulation represents prefrontal disinhibition of supplementary motor areas as movement execution approached, consistent with the established role of right inferior frontal cortex in timing motor inhibition and release (Aron, Robbins, and Poldrack 2004). This reduction in IFG-to-SMA inhibitory connectivity complements the subcortical disinhibition, providing top-down executive permission for motor release.

This integrated pattern of subcortical and cortical activity aligns closely with established neurophysiology of the late CNV. Characterised by a central scalp distribution, the late CNV reflects transition to explicit motor preparation. Our observations of striatal dynamics during late Phase II and Phase III are further supported by intracerebral recordings demonstrating that subcortical preparatory changes in the putamen often precede activity in cortical motor areas (Rektor, Bareš, and Kubová 2001). These results provide strong construct validity for the proposed model, demonstrating its capacity to resolve coordinated subcortical and laminar cortical dynamics underpinning human motor preparation.

##### 3.2.2.2 Motor Inhibition

The mechanisms underpinning motor inhibition during response control engage well-characterised cortical-subcortical circuits, providing a critical testbed for establishing construct validity, specifically the capacity to resolve slow-evolving changes in effective connectivity within established motor control pathways. The presentation of a no-go target demands immediate suppression of a prepotent motor response that engages a distributed fronto-basal ganglia network, in which the rIFG and lpSMA are central to generating the requisite stop command (Aron, Robbins, and Poldrack 2014; Jahfari et al. 2011). Within this architecture, the stop command is relayed via three distinct subcortical pathways that provide nuanced control of motor output. The hyperdirect pathway facilitates rapid global motor inhibition by providing direct cortical excitation of the subthalamic nucleus with exceptionally short conduction times (Nambu, Tokuno, and Takada 2002). Complementing this rapid response, the indirect pathway traversing the cortex, striatum, and GPe to the STN enables more selective suppression of competing motor programmes (Nambu, Tokuno, and Takada 2002; Calabresi et al. 2014). In contrast, the direct pathway facilitates movement through striatal projections to the GPi, effectively disinhibiting the thalamus (Jahfari et al. 2011). Beyond these subcortical routes, the model also accounts for polysynaptic cortical pathways between the lpSMA and lM1 via premotor regions, which exert longer-latency modulatory influences on motor output (Fiori et al. 2016). By applying time-varying parameters to these well-defined connections, we can evaluate whether the model captures the dynamic interplay between executive permission and motor suppression that defines successful response control.

###### Target Processing and Response Selection (Phase IV)

Target presentation initiates a shift in the task set from anticipation toward active response selection, a transition in which the lpSMA appears to function as a primary integrative hub for processing sensory cues and mapping them onto appropriate motor actions (Figure 19). The top-down connection from the rIFG to the lpSMA, which had previously shown reduced coupling during late expectancy, exhibited rapid increase in connectivity upon target presentation (Figure 17). This modulation showed weak support for differential task effects, with further increase for no-go targets (0.18, *p* = 0.78). Simultaneously, the BGT-to-lpSMA connection demonstrated a modest but significant increase (0.44, *p* = 0.93), consistent with evidence that the lpSMA receives dominant subcortical input (Akkal, Dum, and Strick 2007) and serves as an integrative node between frontal response inhibition (Obeso et al. 2013) and subcortical motor planning (Nachev et al. 2007).

**Figure 19.**
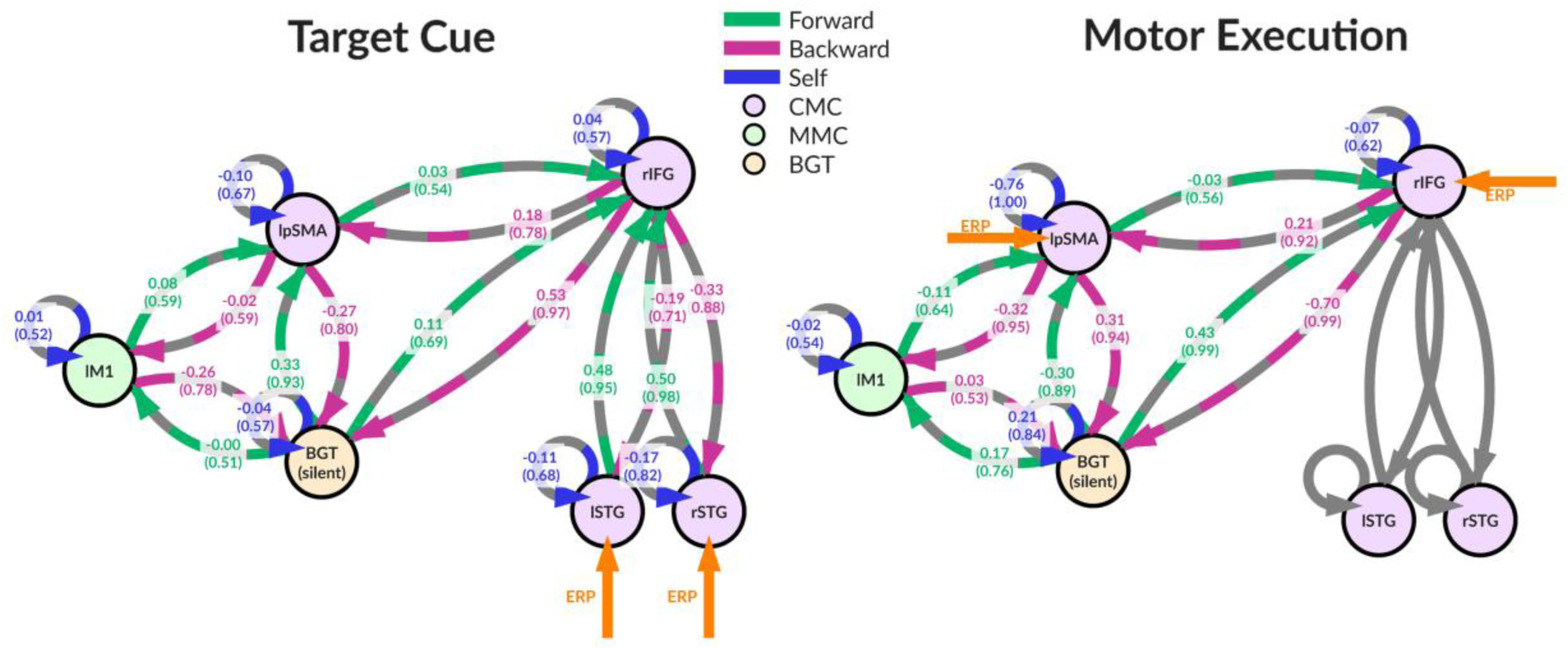
Network connectivity modulation during target cue and motor execution: Empirical modulation of connectivity during target cue and motor execution periods. Network diagrams show posterior expectations for the no-go effect on connection strengths and posterior probabilities (values in brackets) for the target cue (1800 ms) and motor execution (2100 ms) periods. Abbreviations: rIFG, right inferior frontal gyrus; lpSMA, left pre-supplementary motor area; lM1, left primary motor cortex; BGT, basal ganglia-thalamus; lSTG, left superior temporal gyrus; rSTG, right superior temporal gyrus.

Descending connections to subcortical nodes revealed distinct modulation patterns complementing this cortical task set. The projection from the rIFG to the striatum, which initiates both direct and indirect pathways, showed elevated activity across both conditions with further enhancement during no-go trials (0.53, *p* = 0.97; Figure 17). In contrast, the hyperdirect pathway from the rIFG to the STN remained stable near prior values during go trials but demonstrated increased strength during no-go trials, supporting its proposed role in providing rapid motor suppression during response selection to prevent premature execution (Frank 2006). However, because the current model applies a common scaling parameter for experimentally modulated extrinsic connections between any two nodes, trial-specific effects on the hyperdirect pathway cannot be fully disambiguated from those occurring on concurrent striatal pathways. Within the BGT, the hyperdirect pathway (STN to GPi) showed increased connection strength for both conditions, with greater enhancement during no-go trials (Figure 18). Whilst direct and indirect pathways exhibited modest decreases without clear trial differentiation, underlying neural states revealed significant dissociation between striatal and STN activity.

At the faster timescale of neural states, target processing in the frontal regions was characterised by transients in DP firing with the DP depolarisation more marked in rIFG and more profound for no-go trials (Figure 16), suggesting these populations drive the inhibitory signals required to withhold prepotent responses. Striatal populations demonstrated sharp hyperpolarisation across both trial types, with considerable lag during no-go trials (Figure 16). Conversely, the STN exhibited only modest membrane potential variation without clear trial-dependent effects. This dissociation likely reflects fundamental differences in threshold control mechanisms. Striatal activity operates in a bistable regime where phasic responses are shaped by cortical gain modulation (Lee et al. 2019; Bahuguna, Aertsen, and Kumar 2015), whereas the STN maintains tonic pacemaker activity with intrinsically generated bursting properties (Beurrier et al. 1999). These large-amplitude striatal responses may therefore reflect direct cortical drive modulating balance between Up and Down states in medium spiny neurones, whilst modest STN variations reinforce its role as an autonomous pacemaker where threshold control is primarily determined by intrinsic voltage-gated mechanisms (Wilson and Bevan 2011).

###### Response Execution/Inhibition (Phase V)

We now turn to the response execution phase, where connectivity patterns reflected the transition from action selection to motor implementation. The cortico-basal ganglia loops underwent marked reorganisation during motor inhibition (Figure 19). For no-go trials, the connection from rIFG to basal ganglia decreased substantially (−0.70, p = 0.99), whilst the reciprocal connection from basal ganglia-thalamus to rIFG increased (0.43, p = 0.99). This reversal in information flow contrasted with the response selection phase; during motor inhibition, basal ganglia outputs conveyed feedback signals regarding inhibition status back to rIFG, rather than receiving executive commands from the prefrontal cortex (Figure 17). This pattern aligns with the anatomical framework proposed by Haber and colleagues (2003), wherein bidirectional connections between cortex and basal ganglia support both feedforward control and feedback monitoring processes.

The connection from rIFG to lpSMA demonstrated a further increase in coupling strength for no-go trials (0.21, p = 0.92), consistent with sustained inhibitory control demands. The reciprocal connection from lpSMA to rIFG exhibited a sharp increase during this phase, with no differential effect between go and no-go conditions. This absence of task-dependent modulation suggests the pathway operates as a generic monitoring channel, conveying motor status information regardless of whether a response is executed or withheld. Pre-SMA’s connectivity illustrated a distributed pattern supporting motor suppression through both subcortical circuits and direct cortical projections to lM1. For no-go trials, the connection from lpSMA to basal ganglia increased substantially (0.31, p = 0.94), whilst the reciprocal connection from basal ganglia-thalamus to lpSMA decreased correspondingly (−0.30, p = 0.89). Concurrently, the connection from lpSMA to lM1 reduced markedly during response inhibition (−0.32, p = 0.95). These coordinated changes are consistent with pre-SMA’s established role as a crucial node for motor gating (Nachev, Kennard, and Husain 2008), and their detection demonstrates our method’s sensitivity to functionally relevant network reconfigurations.

The BGT-to-lM1 connection representing final output of these basal ganglia pathways did not demonstrate anticipated inhibitory modulation during motor inhibition, instead showing a trend towards increased coupling during no-go trials. Several factors may account for this finding. First, because the BGT node contributes no direct electrophysiological signal, this subcortical component remains less constrained by empirical observations. The model inversion process likely favours cortical parameters more proximal to observed cell populations, potentially reducing sensitivity to subcortical dynamics. Second, alternative architectural configurations may provide superior explanations, as the current model has not undergone formal model comparison. Third, the extended duration of motor execution epochs likely encompasses post-execution evaluative processes, meaning estimated connectivity patterns may reflect motor monitoring rather than execution.

At the faster timescale of neural states, the primary differentiation between go and no-go trials localised to M1. For go trials, DP cells exhibited marked hyperpolarisation entering Phase V, whereas no-go trials showed comparatively modest hyperpolarisation. This finding is counterintuitive, as one would expect substantial depolarisation of DP cells during response execution and hyperpolarisation during response withholding. This may reflect that whilst we applied temporal warping to correct for experimental jitter at target cue onset (as described in Methods), we did not extend this correction to the response phase. Applying response time correction only to go trials would introduce systematic bias between conditions, so we preserved temporal jitter equally across both conditions during this phase. The pronounced hyperpolarisation likely reflects afterhyperpolarisation following corticospinal output [Baldissera et al. (1983); Moberg and Takahashi (2022)], as brief depolarising transients associated with movement initiation would be averaged out across trials, whilst longer-duration afterhyperpolarisations remain prominent in the event-related response. II increased during no-go trials, actively maintaining suppression of DP cells. MP and SP cells showed modest early increases for both conditions, with go trials exhibiting larger transients.

#### 3.2.3 Summary

The empirical analysis demonstrates that the DCM-SR framework successfully tracks plausible neural dynamics across an extended multi-phase task. The framework resolved novel insights into neural generators of the CNV, revealing sustained thalamo-cortical drive targeting prefrontal regions alongside deep-layer hyperpolarisation, patterns consistent with subcortical origins of slow cortical potentials. Across the task sequence, the framework captured coordinated shifts in both neural states and effective connectivity, from engagement of hyperdirect pathways during early motor suppression to progressive disinhibition of thalamo-cortical circuits during late expectancy. During response selection, the model revealed distinct trial-dependent modulations, with enhanced hyperdirect pathway activity and elevated deep-layer firing in frontal regions during no-go trials. One notable limitation concerned the absence of anticipated inhibitory modulation in basal ganglia output to primary motor cortex, likely reflecting that this subcortical node contributes no direct electrophysiological signal and thus remains less tightly constrained during model inversion. Notwithstanding this limitation, the framework has demonstrated the capacity to resolve biologically plausible dynamics at multiple temporal scales within established motor control circuits, providing strong construct validity and establishing a foundation for future model refinements.

## 4 Discussion

The DCM-SR framework addresses longstanding limitations in modelling sequential paradigms, where cognitive processes such as sustained attention and decision accumulation reconfigure network properties between discrete events. It achieves this through three core innovations. First, it generalises non-stationarity across all neural mass parameters, including baseline connectivity, synaptic gains, time constants, and propagation delays. This permits the modelling of fundamental shifts in population dynamics, such as precision-weighted gain control and state-dependent fluctuations in excitability. Second, continuous neuronal state modelling characterises both transient history dependence and persistent path dependence. This facilitates the decomposition of compound signals, such as slow cortical potentials, into evolving synaptic mechanisms, providing a principled approach for inferring the reconfigurations that generate observed responses. Third, the framework formalises transitions as explicit objects of inference through estimable timing, speed, and trajectory parameters, capturing both event-locked exogenous transitions and endogenous shifts. By permitting substantial network reorganisation rather than restricting models to smooth modulation around a reference, DCM-SR is uniquely suited to contexts involving substantive shifts in network architecture.

Validation through simulation and empirical application confirmed that DCM-SR achieves accurate parameter recovery, maintains conservative model selection, and generates plausible neurobiological accounts. Simulations with known-ground truth demonstrated robust recovery across all scenarios, with mean *sRMSE* below 0.36 for modulatory connectivity despite high model dimensionality. Model selection showed an inherent bias towards parsimony, proving more effective at rejecting spurious complexity than detecting genuine variation. For instance, spurious time constants were rejected in 84.4% of cases, while spurious endogenous transitions were rejected in 95.3% of cases. This conservative asymmetry reflects the variational free energy approximation’s tendency to penalise complexity, especially when parameters exert overlapping influences on signal dynamics.

Empirical analysis demonstrated the framework’s ability to resolve dynamics across extended multi-phase paradigms. By tracking concurrent neural state evolution and connectivity reconfigurations, DCM-SR provided the first biophysical model of human CNV generation. The framework decomposed the CNV into evolving thalamo-cortical drive, laminar-specific membrane dynamics, and shifting basal ganglia engagement. These results revealed that sustained subcortical input and deep-layer hyperpolarisation, rather than superficial depolarisation, drive the CNV. Additionally, the model captured trial-dependent hyperdirect pathway strengthening during motor inhibition. By separating overlapping processes operating at different timescales, DCM-SR offers a mechanistic window into complex human cognitive dynamics previously accessible only through invasive recordings.

### 4.1 Wider Applications in Cognitive Neuroscience

The integration of expressive, time-varying network reconfigurations with continuous neural dynamics facilitates new mechanistic investigations into the temporal architecture of cognition. DCM-SR’s capacity to track evolving parameters is uniquely suited to examining processes where network properties reconfigure over behaviourally relevant timescales. This includes attentional control systems, where shifting priorities require the dynamic regulation of network gain and selectivity (Corbetta and Shulman 2002; Petersen and Posner 2012), and neuromodulatory systems, where acetylcholine, dopamine, and noradrenaline alter synaptic efficacies and integration windows according to arousal and task demands (Marder 2012; Doya 2002).

Furthermore, the framework accommodates metacognitive monitoring, which requires sustained adjustments to operations across extended sequences (Fleming, Dolan, and Frith 2012), and perception-action loops, where sensorimotor contingencies evolve through environmental interaction (Cisek and Kalaska 2010). We focus on two domains where substantial theoretical development exists regarding dynamical systems properties, yet translation to human non-invasive neuroimaging remains limited. Decision-making and planning have been extensively characterised through attractor landscapes, regime transitions, and bifurcations in animal models (S. Wang et al. 2023; Thura et al. 2022; Kaufman et al. 2014), whilst working memory exhibits fundamental history dependence and hysteresis, with neural responses depending on the accumulated trajectories of preceding events (Barbosa et al. 2020; Lundqvist et al. 2016). These domains exemplify processes where identifying the generative mechanisms of time-varying parameters and path-dependent evolution is essential for bridging computational theories with biophysical implementations in human neural circuits.

#### 4.1.1 Decision Making and Planning

Decision making involves evaluating counterfactuals based on sensory evidence, while planning extends this to action sequences directed toward goal states. These multi-phase processes operate across distinct timescales (Cochrane et al. 2023): rapid sensory encoding occurs over tens of milliseconds, intermediate evidence accumulation unfolds over hundreds of milliseconds to seconds as populations integrate information toward commitment thresholds (Ratcliff and McKoon 2008; Evans et al. 2017), and strategic adjustments or learning occur over minutes to days. These temporally diverse processes are implemented by neural populations exhibiting dynamics ranging from fast transients to slow baseline drift.

Research in rodents and non-human primates characterises decision making as transitions between dynamical regimes within attractor landscapes (Boyd-Meredith et al. 2022; S. Wang et al. 2023). During deliberation, neural trajectories evolve within low-dimensional manifolds shaped by evidence and urgency before undergoing rapid phase transitions at commitment, shifting activity into orthogonal subspaces for motor preparation (Thura et al. 2022; Kaufman et al. 2014). These transitions often reflect bifurcations, where crossing critical boundaries triggers qualitative network reorganisation rather than gradual change (Albantakis and Deco 2011; X.-J. Wang 2012). Timescale separation emerges as fast sensory dynamics operate within attractor basins that evolve slowly through learning, with attractor stability directly reflecting decision confidence (S. Wang et al. 2023; Tang, Shin, and Jadhav 2021).

Despite these insights from invasive recordings, human neuroscience has largely remained limited to correlating activity with cognitive model parameters like drift rates and thresholds (Forstmann et al. 2008; Mulder, van Maanen, and Forstmann 2014). While such approaches identify regions tracking decision variables, they lack mechanistic accounts of how neuronal processes generate these dynamics. DCM-SR bridges this gap by linking decision phenomenology to biophysical mechanisms, characterising how synaptic efficacies and time constants reconfigure to implement processing phases like accumulation and urgency-driven commitment. The framework can test whether activity reflects smooth parameter drift or abrupt regime shifts analogous to bifurcations. Crucially, it accommodates the hierarchical timescales of decision making by estimating the timing and nature of reconfigurations, providing a principled method for connecting cognitive theories with the biophysical properties of human neural circuits.

#### 4.1.2 Working Memory and Maintenance

Working memory supports the active maintenance and manipulation of information, serving as a cognitive workspace for reasoning and goal-directed behaviour. Traditional accounts attribute maintenance to persistent activity in recurrent networks, where sustained firing maintains information in stable attractor states (Compte et al. 2000; X.-J. Wang 2001). This view is challenged by activity-silent models proposing that short-term synaptic plasticity, such as calcium-mediated facilitation, maintains information without continuous spiking (Mongillo, Barak, and Tsodyks 2008; Stokes 2015). Recent evidence suggests these mechanisms interact dynamically; attractor dynamics and activity-silent substrates coexist in the prefrontal cortex, with their relative contributions shifting based on task demands (Barbosa et al. 2020; Murray et al. 2017) and whether information is passively maintained or actively manipulated (Trübutschek et al. 2019). Animal electrophysiology suggests this interplay involves rapid, context-dependent modulation of recurrent excitation, inhibitory gain, and NMDA-mediated slow dynamics. These processes span multiple timescales, from fast gamma oscillations to slow attractor drift unfolding over seconds (Lundqvist et al. 2016; Barbosa et al. 2020). DCM-SR can readily capture these dynamics by allowing synaptic parameters to evolve across sequential responses, accommodating slow NMDA kinetics alongside faster AMPA-mediated excitation and trial-by-trial adaptive changes in inhibitory gain.

Capturing time-varying parameters alone, whilst necessary, remains insufficient to fully characterise working memory dynamics. Working memory exhibits fundamental history dependence, where the neural response to a stimulus depends not only on the current input but on the accumulated trajectory of preceding events (Barbosa et al. 2020). This manifests behaviourally as serial biases, where information from one trial influences the next. Computationally, these effects arise from the interaction between attractor dynamics, which structure the phase space, and activity-dependent plasticity, which modulates the landscape over time. The critical advance of DCM-SR lies in modelling this interplay explicitly. By allowing parameters to evolve continuously while representing neural states as trajectories through state space, the framework captures genuine hysteresis. System response properties change as parameters drift, meaning a previously encountered stimulus produces a different response because the underlying synaptic landscape has been reshaped. This path-dependent evolution, where parameter changes accumulate irreversibly, distinguishes working memory from simple recurrent dynamics. Without accounting for such hysteresis, models risk treating each response as arising from a reset system, misrepresenting how synaptic strengths drift under cognitive demand. DCM-SR provides the machinery to disambiguate slow synaptic dynamics from path-dependent hysteresis, offering insights into how the human brain implements flexible information storage over behaviourally relevant timescales.

### 4.2 Conclusion

The development of DCM-SR provides a biophysically grounded account of how network configurations evolve alongside the neural states they support. Our validation confirms that this increased model complexity remains parsimonious, with robust parameter recovery and conservative selection properties ensuring that inferred reconfigurations reflect genuine dynamical shifts rather than statistical overfitting. The empirical decomposition of the contingent negative variation demonstrates the utility of this approach, revealing a sophisticated interplay of subcortical drive and laminar-specific cortical responses that eludes traditional signal-averaging methods. Ultimately, the capacity to resolve hierarchical timescales and path-dependent evolution establishes a principled bridge between the computational rigour of animal electrophysiology and the non-invasive observation of human cognition. This provides a necessary foundation for investigating how the human brain implements the extended temporal structures essential for complex, goal-directed behaviour.

